# Polybacterial intracellular coinfection of epithelial stem cells in periodontitis

**DOI:** 10.1101/2023.08.23.554343

**Authors:** Quinn T. Easter, Bruno Fernandes Matuck, Germán Beldorati Stark, Catherine L. Worth, Alexander V. Predeus, Brayon Fremin, Khoa Huynh, Vaishnavi Ranganathan, Diana Pereira, Theresa Weaver, Kathryn Miller, Paola Perez, Akira Hasuike, Zhaoxu Chen, Mandy Bush, Blake M. Warner, Janice Lee, Shannon M. Wallet, Inês Sequeira, Katarzyna M. Tyc, Jinze Liu, Kang I. Ko, Sarah A. Teichmann, Kevin M. Byrd

## Abstract

Periodontitis affects billions of people worldwide. To address interkingdom relationships of microbes and niche on periodontitis, we generated the first sin-gle-cell meta-atlas of human periodontium (34-sample, 105918-cell), harmonizing 32 annotations across 4 studies^1–4^. Highly multiplexed immunofluorescence (32-antibody; 113910-cell) revealed spatial innate and adaptive immune foci segregation around tooth-adjacent epithelial cells. Sulcular and junctional keratinocytes (SK/JKs) within epithelia skewed toward proinflammatory phenotypes; diseased JK stem/progenitors displayed altered differentiation states and chemotactic cytokines for innate immune cells. Single-cell metagenomics utilizing unmapped reads revealed 37 bacterial species. *16S* and rRNA probes detected polybacterial intracellular pathogenesis (“co-infection”) of 4 species within single cells for the first time in vivo. Challenging coinfected primary human SK/JKs with lipopolysaccharide revealed solitary and synergistic ef-fects. Coinfected single-cell analysis independently displayed proinflammatory phenotypes in situ. Here, we demonstrate the first evidence of polybacterial intracellular pathogenesis in human tissues and cells—potentially influencing chronic diseases at distant sites.

## INTRODUCTION

Periodontal diseases—i.e., periodontitis—affect billions globally every year^5^. They are characterized by dysregulated, chronic periodontium in-flammation—most often caused by polybacterial dysbiosis^6^; if left untreated, the result is tooth loss^7^. Precision medicine approaches for periodonti-tis—including diagnostics, prognostics, and biologics—have had minimal success to date^8,9^. Periodontal diseases are associated with >60 systemic diseases; early identification and treatment may improve overall health^10^. A limited understanding of cell subpopulations and their cell states, either supporting niche maintenance or contributing to its breakdown, inhibits precision approaches. There is an unmet need to elucidate cell-specific and cooperative cell plasticity in periopathogenesis. Despite twenty-five-year knowledge of tooth-associated keratinocyte heterogeneity^11^, cell type functional annotation remains incomplete—especially considering mono/ polybacterial infection.

Structural immunity^12^ i.e., “immune functions of non-hematopoietic cells”, can be studied using single-cell and spatial genomic approaches. Recent work across disciplines now suggests that in certain contexts “every cell is an immune cell”^13^. In periodontitis, this concept has been demonstrated between fibroblasts and innate immune cells^4,14^. Other structural cell types such as epithelial cells—specifically keratinocytes—have been implicated in regulating autoimmune and infectious disease immune responses^15–17^. Furthermore, while epithelial cells/keratinocytes may passively permit microbial infection via tissue barrier breakdown^18^, keratinocytes via their stem/progenitor cells may play important immune education roles at the tissues barrier before, during, and after barrier breakdown; alternatively, keratinocyte stem/progenitor cell rewiring may retain the capacity for unique response to chronic challenge^19,20^.

To address outstanding tooth-associated epithelia structural immune roles, we created an integrated periodontitis meta-atlas of human tissues^1–4^ using an open-source single-cell analysis toolkit (Cellenics®; https://github. com/hms-dbmi-cellenics), describing 17 total and functionally annotating 4 new gingival keratinocyte subpopulations within the human gingival epithelium at a single-cell level: tooth-facing sulcular keratinocyte (SK) and tooth-interfacing junctional keratinocyte (JK) stem/progenitor cells and their differentiated progeny were marked by KRT19 expression. We validated these cell identities using multiplexed in situ hybridization (mISH); mice lack evidence for Krt19 tooth-associated keratinocytes. Using highly multiplexed immunofluorescence assays (mIF; 32-antibody), we show immune foci differences between these SK and JK niches, including more immune suppression phenotypes at areas closest to the tooth in disease. Transcriptionally, SK and JK microniches respond differently in disease but generally alter cell differentiation and upregulate effector cytokines (“epi-kines/keratokines”) along stem-cell-to-progeny trajectories—predominantly in JKs.

We also predicted these cells differentially regulate innate and adaptive immune cell subpopulations, even in health. We combined unmapped reads of our integrated atlas, mIF, and mISH and found that some SKs and JKs tolerate intracellular pathogenesis (i.e., polybacterial coinfection, PIC) of numerous periodontal pathogens in epithelial stem cells and their progeny—the first example of at least 4 distinct intracellular species within single cells. PIC models of human gingival keratinocytes in vitro confirmed upregulated keratokine signatures with and without extracellular lipopolysaccharide. mISH in vivo revealed cell-specific PIC signatures mirroring in vitro models, revealing regional phenotypes mostly associated with 16S signal within tissue microniches. Here, we demonstrate the first evidence of polybacterial intracellular pathogenesis in human tissues and cells driving epithelial stem/progenitors and their progeny toward proinflammatory cell states, potentially accumulating in healthy tissues to influence chronic diseases over the lifespan.

## RESULTS

### Generation and analysis of a first draft integrated periodontitis meta-atlas

The tooth is supported by diverse cell types (Figure 1a). Hundreds of diseases affect teeth; however, the cell-specific contribution to these diseases remains limitedly explored^21^. Recent murine^22–24^ and human^1–4^ studies have focused on the single-cell RNA sequencing (scRNAseq) of the tooth-supporting periodontium (mineralized: alveolar bone, cementum; soft: gingiva, periodontal ligament)^26^, but some cell types like keratinocytes are minimally annotated despite knowledge of their heterogeneity^27^. We analyzed 4 human scRNAseq datasets^1–4^ (34 samples, 3 states; Figure 1b) using Cellenics®. Metadata was harmonized (Supplementary Table 1)^28^ and the location noted for 27/34 samples to establish a common coordinate framework (CCF) for periodontium^29^ (Extended Data 1a).

**Figure 1.**
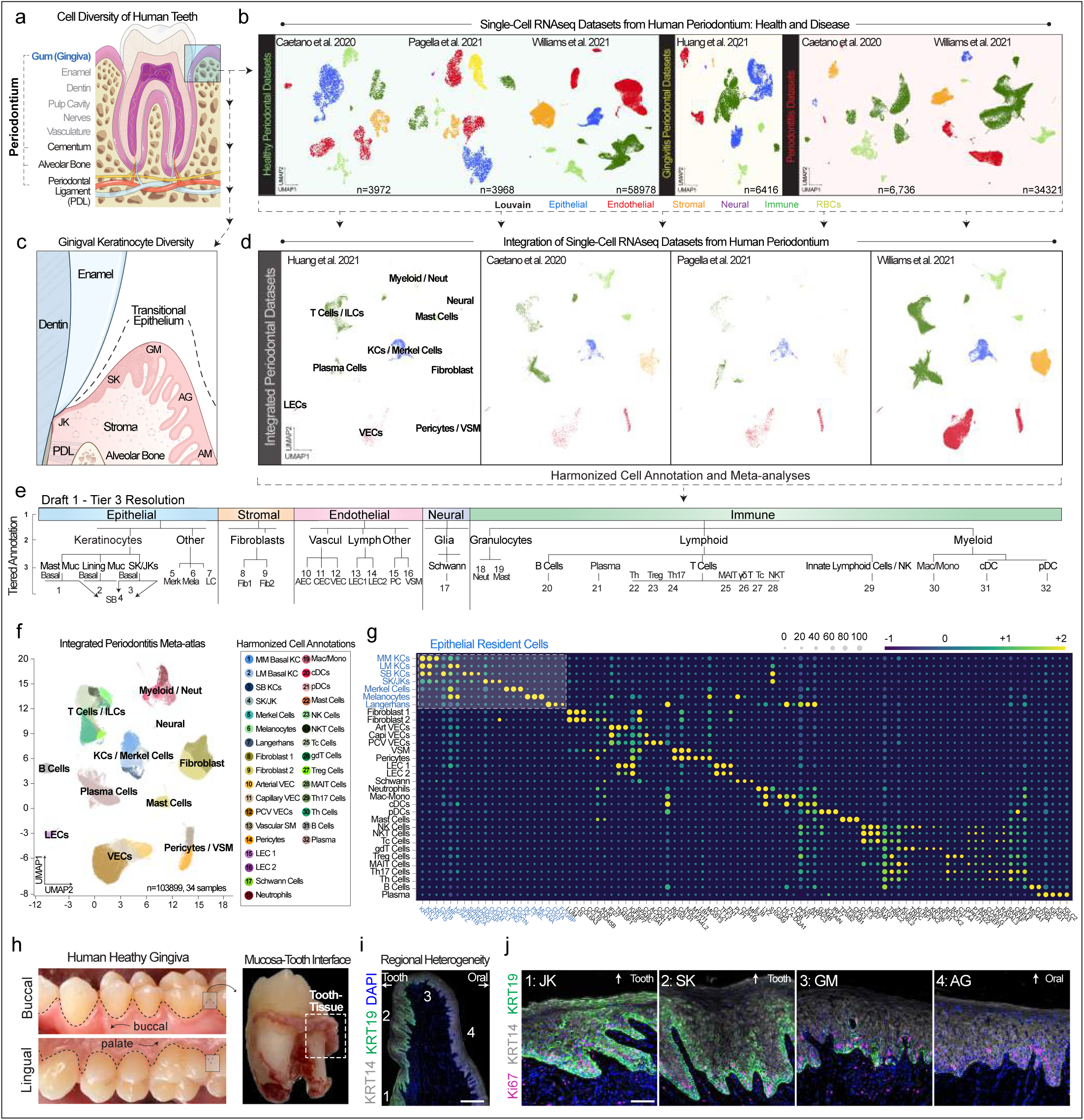
Creation of the first draft of an integrated periodontitis meta-atlas. (a-e) (a) Human teeth are supported by a diverse collection of specialized tissues, including the periodontium consisting of the gingiva (blue: epithelial and stromal tissues), periodontal ligament, and mineralized tissues (cementum and alveolar bone). (b) Four studies to date have profiled the soft tissue components of human periodontium in health and disease states^1–4^. Datasets were first reprocessed and using the Biomage-hosted community instance of Cellenics®, broad cell classes were compared between the four studies, underscoring the complementary nature of cell capture based on unique dissociation strategies among the studies. (c) It is known that the epithelial attachment of the gingiva is also specialized but remains poorly understood. This niche is a stark example of transitional epithelia, and depending on the site, changes from non-keratinized alveolar mucosa (AM), to keratinized attached gingiva (AG), begins to alter expression profiles at the gingival margin (GM) before specializing in the gingival sulcus epithelial keratinocytes (SK) and junctional epithelial keratinocytes (JK). (d) Each study was first integrated using Harmony and assigned Tier 1 cell type annotation. (e) Each dataset was further integrated and harmonized tier annotation was performed between epithelial, stromal, endothelial, neural, and immune cell populations. No single study fully represented each cell type. (f-h) Integrated UMAP, cell assignments (f), and cell signatures (g; Supplementary Table 1) were generated for the meta-atlas; epithelial cells (blue) are highlighted in the shaded box. For public use, the entire dataset was uploaded to cellxgene. (h-j) SK and JKs were grouped in the Tier 3 analysis as co-expressing Keratin 14 (*KRT14*) and Keratin 19 (*KRT19*); Supplementary Table 1. (h) To validate this, healthy human gingival tissues were preserved on the tooth surface after extraction and fixed. (i) IHC reveals the gradual transition from tooth-facing JK and SKs to GM, AG, and AM keratinocytes using KRT19; however, (j) KRT19-high and -low epithelial stem cells proliferate in the basal layer in health. Abbreviations: Innate Lymphoid Cells (ILCs); Keratinocytes (KCs); Vascular Endothelial Cells (VECs); Vascular Smooth Muscle (VSM); Lymphatic Endothelial Cells (LECs); Neutrophils (Neut); Masticatory (Keratinized) Mucosa (Mast Muc; MM); Lining (Non-Keratinized Mucosa (Lining Muc; LM); Suprabasal (Differentiated) Keratinocytes (SB); Fibrolast (Fib); Arterial Endothelial Cells (AECs); Postcapillary Venule (PCV); Venule Endothelial Cells (VECs); Macrophage/Monocytes (Mac/Mono); Conventional Dendritic Cells (cDCs); Plasmacytoid Dendritic Cells (pDC); Cytotoxic T Cells (Tc); Gamma Delta T Cells (gdT); Regulatory T Cells (Treg); Mucosal Associated Invariant T Cells (MAIT); Helper T cells (Th). Illustration from (a) created with https://bioicons.com; illustration from (b) created with BioRender.com. Scale bars: 250 and 50 μm.

All samples were reprocessed, filtered, and integrated (Figure 1b; see Methods). Cells were broadly annotated (Tier 1 resolution). We focused on gingival epithelial heterogeneity (Figure 1c) within the distinct transitional zone between non-keratinized alveolar mucosal (AM), attached gingival (AG), gingival margin (GM), and sulcular and junctional keratinocytes (SK/JKs)27. Red blood cells were filtered; each study was integrated and further annotated (Tier 2; Figure 1d). Integrating data enabled the harmonized cell annotation of 32 cell types across datasets (Figure 1e,f). Epithelial cells could be classified into 7 different types, including SK/JKs. Comparing each study, cell type proportions revealed complementary subpopulations (Extended Data 1b). Cellenics® data was exported to cellxgene for public use (Extended Data 1c).

Marker genes were determined for each of the 32 cell types (Supplemen-tary Table 1). Keratinocytes were broadly marked by *KRT14/KRT5*. SK/JKs expressed higher *FDCSP*, *ODAM*, and baseline interleukin/chemok-ine expression, suggesting active roles in inflammation by these tooth-as-sociated keratinocytes. One significantly upregulated marker in SK/JKs was another keratin, *K19*/KRT19^11^. To validate the K19/KRT19 spatial localization, adult gingiva was harvested, and orientation was preserved to feature both oral-facing and tooth-facing keratinocytes (Figure 1h). Immu-nohistochemistry validated K19 as the definitive SK/JK marker (Figure 1i). Each of these regions within the entire gingiva revealed similar proportions of Ki67+ cycling cells, highlighting the need to understand SK/JK epithelial stem/progenitor cells in humans, similar to mice^30^ (Figure 1j).

### Spatial proteomic analysis of periodontitis reveals distinct peri-epithelial immune foci

Research into periodontitis inherently has a spatial dimension due to oral and tooth-facing tissue polarity. This is relevant because facultative and obligate aerobes predominate within biofilms on the tooth and mucosal surfaces nearest the tooth-soft tissue attachment (i.e., junction; Figure 1a,c, 2a)--especially in disease states. Informed by our meta-atlas, we designed a highly multiplexed immunofluorescence (mIF) assay (32-antibody; 113,910-cell) across healthy and periodontitis samples to understand how disease affects spatial cell arrangements (Figure 2b, Extended Data 1c,d; Supplementary Table 1). In periodontitis, we consistently found concentrated CD45+ adaptive immune cells near SK cells; we also found isolated expression of KRT19 cells in the keratinized mucosa (attached gingiva) uniquely attracting CD45+ immune cells (inset, Figure 2b), suggesting KRT19+ cells may be inherently chemotactic.

**Figure 2.**
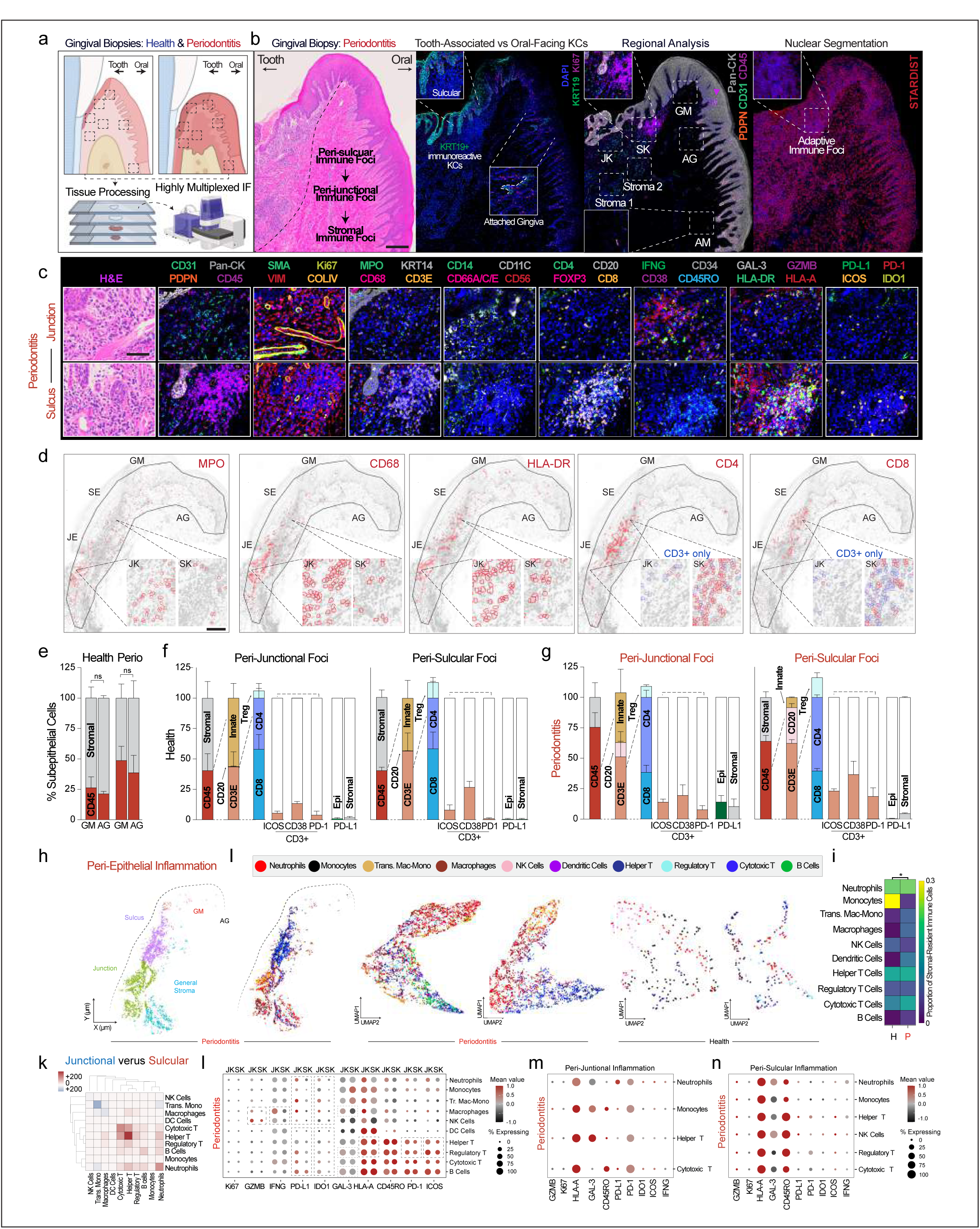
Highly multiplexed, spatial immunofluorescence assays of periodontitis reveal innate and adaptive immune cell foci and phenotypes in peri-epithelial niches. (a-c) (a) The orientation of periodontal tissues is critical to show a tooth-facing (junctional, sulcular epithelial keratinocytes; SK/JKs) and oral-facing (gingival margin, GM; attached gingiva, AG; and alveolar mucosal, AM keratinocytes) attachment for highly multiplexed immunofluorescence (mIF) assays of periodontitis. (b) By doing this in sequential sections, we first confirmed orientation and noted highly localized inflammatory profiles near tooth-facing epithelial keratinocytes. As discovered in the initial analysis (Figure 1), tooth-facing JK and SKs uniquely express Keratin 19 (KRT19) in every cell type, highlighting the transition zone. An initial analysis of Tier 1 cell assignments using mIF (PhenoCycler-Fusion (Akoya Biosciences) revealed adaptive immune foci concentrated near SKs and more diverse, innate immune populated foci near JKs. Cell segmentation was performed using StarDist. (c) Expanding the assay to include 32 antibodies revealed more heterogeneity at the cell type and cell state level. Antibodies are grouped and zoomed-in regions from a periodontitis sample is featured; more regions from (b) are also shown in Extended Data 2d. (d-g) (d) Manual thresholding was performed to show individual marker heterogeneity along the peri-epithelial niches, Cell assignment algorithm for this study is shown in Extended Data 2. (e) Single marker analysis of CD45+ cells revealed general increases in both GM and AG stroma. (f) Comparison of cell identities and cell states shows minimal difference between peri-junctional and peri-sulcular niches in health. (g) While both peri-junctional and peri-sulcular immune infiltrate increase in disease, peri-junctional foci are biased toward more innate and relatively more CD4+ T cells compared to peri-sulcular niches, which are biased toward adaptive immune cells (T and B Cells). Both junctional stromal and JKs also express more PD-L1 compared to sulcular cell types. (h-k) (h) Spatial analysis of peri-epithelial regions was broken into four specific and one broad classification. (i) Segmented immune cells were assigned identities in health and disease. (j) Periodontitis displays more diverse heterogeneity considering the whole tissue. (k) However, cell-cell interactions among immune cells revealed diverse enrichment of immune cell types in peri-junctional and peri-sulcular immune foci in periodontitis. (l-n) Cell states are also spatially distinct in periodontitis, with the peri-junctional immune cells expressing more immune exhaustion. Abbreviations: Antibodies (see Methods); also see Figure 1 legend. Scale bars: 250 and 50 μm. Statistical test (e): p<0.05, paired Student’s T-test (e); chi-square test (j). Illustration from (a) created with BioRender.com.

For whole-slide analysis, we segmented images using StarDist (Figure 2b, see Methods). The junctional region consistently revealed higher innate immune cell concentrations (MPO+-Neutrophils, CD14/CD68+ Macrophages, CD56+-Natural Killer cells, CD11c+-Dendritic cells); the sulcular region revealed distinct adaptive immune foci (CD8+-Cytotoxic T Cells, CD4+-Helper T Cells, FOXP3+-Regulatory T Cells, and CD20+-B Cells). (Figure 2c,d; Extended Data 1e). We quantified this by region using single markers and manual thresholding, revealing more infiltrate in peri-epithelial stroma (Figure 2e) and higher innate peri-junctional immune foci frequency in disease (Figure 2f,g). This extended to cell states of CD3+-T Cells, which displayed more ICOS+, CD38+, and PD 1+-T cells in peri-sulcular foci (Figure 2g). The proportion of PD-L1+ cells was nearly 10x higher in health and 2x in disease in the peri-junctional stroma, yet nearly 3x higher in health and 100x higher in disease when analyzing JKs compared to SKs (Figure 2g; Extended Data 1f,g). Thus, the periodontal niche may potentially support immunosuppressive microenvironments nearest to the tooth-soft tissue interface, but the reason for this was unknown.

Using multiple protein markers, cells were assigned tiered identities (Figure 2d; Extended Data 1h, see Methods) considering tooth proximity (Figure 2h,i). Each peri-epithelial immune foci immune constituent was assigned an identity in these regions of interest (ROIs). Proportionally, tissue-wide, immune cell ratios shifted to favor dendritic, macrophage, cytotoxic T, and B cells (Figure 2j). Considering local neighborhoods, the sulcus supported more immune-immune predicted “interactions” within cellular neighborhoods, favoring both innate and adaptive immune cell types; however, the junction supported interactions between CD68/CD14+ transitioning monocytes/macrophages, CD68+ macrophages, and MPO+ neutrophils, highlighting the chronic peri-junctional immunosuppressive niche impact, which validated manual thresholding approaches (Figure 2k). Assessing cell states, peri-junctional immune cells expressed more GZMB, IFN-γ, Galectin-3, PD-L1, and HLA-A in disease compared to peri-sulcular foci (Figure 2l-n). Overall, the junctional zone structural niche appeared to be more immunosuppressive than other peri-epithelial niches, but the underlying reason remained unclear.

### Transcriptomic analysis reveals immune roles for and effects on keratinocytes in periodontitis

Recent work supports the idea that “structural” cells (i.e., vasculature, fibroblasts, epithelial cells) can coordinate to tailor individual immune responses to niche-specific challenges (“structural immunity”^12^). In periodontitis, this has been relatively unexplored. Individually, fibroblasts in single-cell^4^ and spatial transcriptomic^14^ studies have shown direct innate immune curation roles. To ask how structural cell types may play roles in periodontitis progression, we analyzed periodontitis versus healthy cells in a pseudobulk RNAseq analysis, generating differentially expressed gene (DEG) lists (Extended Data 2a, Supplementary Table 1) for keratinocytes, fibroblasts, and vasculature endothelial cells (VECs; Extended Data 2b-d). We observed structural populations comprising ∼2/3 of the up-and down-regulated DEGs from the pseudobulk experiment (Extended Data 2e), supporting structural immune roles in periodontitis.

Considering the unique localization of immune cells in peri-epithelial niches, we wanted to understand how keratinocytes are altered in periodontitis. Our analysis revealed upregulation of *CXCL1, CXCL3, CXCL8, CXCL13, CCL20, CSF3, IL1A, IL1B*, and *IL36G* and receptors *IL1R1*, *IL7R* in periodontitis; furthermore, *CXCL1, CXCL8, IL1A,* and *IL1B* had a greater log fold change in keratinocytes compared to all cells (Extended Data 2), and *CXCL17, CCL20*, and *CSF3* were uniquely upregulated in keratinocytes. We used g:Profiler (see Methods) to understand altered disease state pathways, using functional profiling of GO pathways to highlight cell signaling and bacteria/dysbiosis responses as primary up-regulated responses. (Extended Data 3a). Immune response upregulation likely comes at the expense of terminal differentiation, development, and translation (Extended Data 3b). This phenomenon, in concert with stromal populations, appears to reshape the periodontal niche primary via immune education.

Using CellPhoneDB (see Methods), we investigated relative cell-cell communication patterns of Tier 3-level cell identities via receptor-ligand pairs (Extended Data 3c, Supplementary Table 2). Complementing expected fibroblast-vasculature and fibroblast-immune interactions, we found significant predicted communication between keratinocytes and many other cell types, suggesting regional epithelia-stromal resident population communication in periodontitis—most prominent for SK/JKs compared to other epithelial cells (keratinocytes, melanocytes, Merkel cells). Even in health, SK/JKs were enriched for cytokines and matrix metalloproteinases (Extended Data 3d)—relatively higher compared to other structural cell types. In health and disease, other stromal cells upregulated cytokines complementary to SK/JKs (Extended Data 3d). SK/JKs were robustly active in health when analyzing receptor-ligand pairs (Extended Data 3e). The immunoregulatory role of keratinocytes in periodontitis has been studied^18,31^, and the innate immune population residency near JKs has been shown^32^; however, while the “reactome” of diseased keratinocytes further suggests active immune roles for these cells (Extended Data 3f), direct roles for SK/JK epithelial stem/progenitors and their progeny remain understudied.

### Redefining human gingival keratinocyte subpopulations for niche-specific analysis in periodontitis

Considering both the single-cell and spatial datasets, we suspected SK/ JKs might represent new human cell types. We subclustered keratinocytes from our integrated atlas (∼8500 cells, Extended Data 4a, 5a) and generated new markers for each population (Extended Data 4b, Supple-mentary Table 1). We validated a robust *KRT19*-high population uniquely clustering within the dataset. Using a custom 12-plex in situ hybridization (ISH) assay (RNAscope) designed from single-cell signatures with built-in negative/low controls, we refined cell cluster annotations, finding opposite *CXCL14* presentation to KRT19 IHC (Figure 1i,j) and enrichment in AG and the GM transitional zone (Extended Data 4c, 5b). Other markers enriched in oral-facing keratinocytes included *NPPC, PAPPA*, and *NEAT1* but *SAA1, IL18*, and *RHCG* for SK/JKs. Using primary human gingival keratinocytes (HGKs), we discovered the presence (Extended Data 4d) and maintenance (Figure 4e,f) of KRT19+ cells. Using ISH, we also found cell subpopulation-specific markers such as *FDCSP*—defining SK/JKs (Figure 1; Extended Data 4g). This established multiple lines of evidence showing SK/JK epithelial stem cells and their progeny in vivo and models for understanding their cell subpopulation-specific activity and disease response.

### Murine gingival keratinocyte subpopulations exist around molars but are less diverse

Despite frequent use of mouse as a periodontitis model^33^, the junctional niche similarity between mice and humans is debated^34,35^. To address this, we performed scRNAseq of adult mouse gingiva (Extended Data 5c). Subclustering epithelial cells revealed a subpopulation of Krt19-high keratinocytes. When compared to basal epithelial stem/progenitors (Krt14+) and differentiated (*Krt4+, Krt1+*) keratinocytes, *Krt19+*-cells were mostly *Krt14+* (Extended Data 5d). IHC analysis of healthy mouse gingiva around molars (M2; M3) readily showed *Krt14* and *Ki67* but not *Krt19*, suggesting no gene-to-protein translation (Extended Data 5e). These rare *Krt19* cells also upregulated *Cxcl1, Cxcl8*, and *Cxcl17*, distinct from other *Krt14*- high cells (yellow box: Extended Data 5f), suggesting some mirroring of human proinflammatory cells; yet some other mixed cell subpopulations had matched interleukin expression (pink box: *Il1a, il1b, Il1rn, Il18*). When using the same human signatures, mouse subpopulations failed to segregate, supporting only some heterogeneity of murine gingival keratinocytes, warranting further investigation of species-species cell type annotations^36^.

### Periodontitis affects SK/JK stem/progenitor differentiation to upregulate inflammatory signatures

Our confirmation of SK/JK cell identities led us to refine cell annotations using new markers (Figure 3a) and consider how SK/JKs are affected in disease. To better understand SK/JK heterogeneity, we subclustered out the KRT19-high cells from the integrated atlas, annotating 17 keratinocyte populations (Figure 3b,c) to include basal (stem/progenitors) and their differentiated progeny (suprabasal keratinocytes; SB). We next looked at the cell-specific gene upregulation patterns to gain insight into these cells in periodontitis (Supplementary Table 1). Despite their spatial adjacency, we found 28.5% of shared SK/JK gene upregulation in periodontitis (Figure 3d). We found basal and SB JKs—not basal and SB SKs—generally upregulated effector cytokines *CXCL1, CXCL6, CXCL8, IL1B*, and *IL36G* (*CXCL1, IL1B* by SB JKs compared to SB SKs (Figure 3e)).

**Figure 3.**
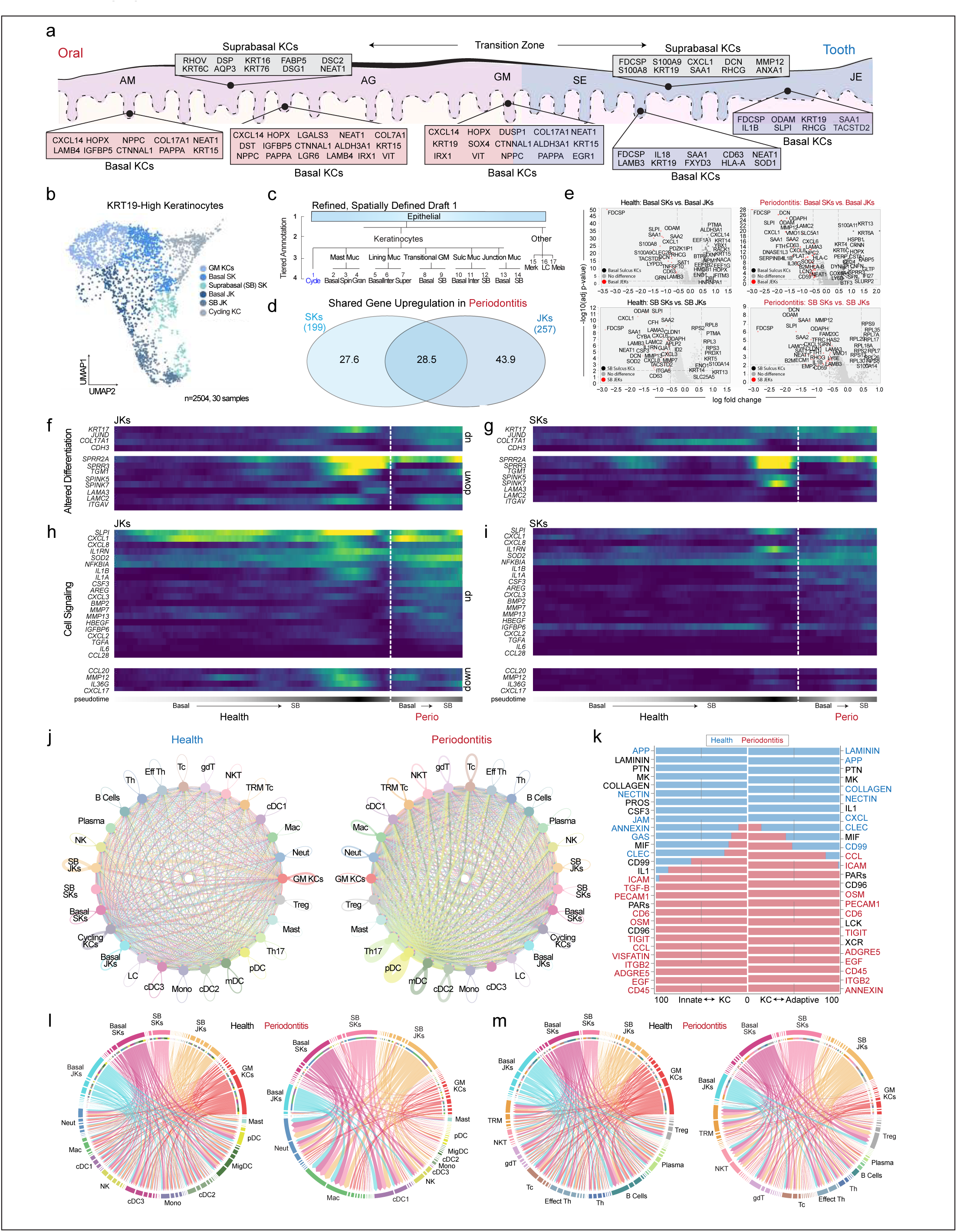
Proinflammatory profiles coincide with altered differentiation trajectories of tooth-associated epithelial stem/progenitor cells. (a-c) (a) Due to the single-cell annotation and in situ validation, a draft model of the oral-to-tooth transition zone in humans is presented with basal and suprabasal keratinocyte markers. (b) These markers allowed for *KRT19*-high keratinocyte (KCs) cell subclustering for the first time (2504 cells in total), including gingival margin keratinocytes (GM), sulcular keratinocytes (SK), and junctional keratinocytes (JK). (c). This also permitted a more granular draft annotation of cells of the gingival attachment. (d-e) (d) Assaying differentially expressed genes in periodontitis, SK and JKs only share about a quarter of upregulated genes, further underscoring spatial differences in their response in vivo. JKs display nearly 125 unique upregulated genes in diseased cells. (e) Further analysis considering basal versus suprabasal (differentiating) keratinocytes reveals unique cell signatures between basal and suprabasal cell types. This full list is included in Supplementary Data 1. (f-i) To understand SK and JK trajectories, we used partitioned-based graph abstraction (PAGA) on these cell types, comparing these cells between basal and suprabasal cells and in health and disease states using pseudotime (f) Looking at the basal to suprabasal transition, aligned cells from JKs display altered gene expressed comparing health to disease cell types including broader expression of *KRT17*, more expression of *JUND*, *COL17A1*, and *CDH3*. Key differentiation genes such as *SPRR* family members are downregulated. (g) JKs display robust cell signaling and inflammatory phenotypes, which are exacerbated in disease states. (h). In SKs, differentiation genes are uniquely expressed compared to JKs in health but also appear altered in disease along basal to suprabasal trajectory. (i) SKs appear generally less reactive compared to JKs in disease. (j) CellChat was used to understand cell signaling pathways in health and disease considering tooth-associated keratinocytes. Circle plots highlight the significant receptor-ligand interactions between any cell populations, including same-cell type signaling interactions (i.e., JK-JK, SK-SK, etc.). The proportion of interactions increased across more detailed immune cell type annotations (Tier 4 annotations; see Extended Data 7). (k) Relative information flow in health and disease shows a preference for cell adhesion (NECTIN, COLLAGEN, JAM, LAMININ) and other pathways such as APP, CXCL, and MIF pathways. In disease, more preference for cell signaling pathways is preferred, such as TGFB, TIGIT, CCL, CD45, and EGF. Innate (l) and adaptive (m) immune cell communication was measured gene by gene using a chord diagram for visualizing cell-cell communication. Abbreviations: Cycling Keratinocytes (Cycle; KCs); Spinous Layer (Spin); Granular Layer (Granular); Intermediate Layer (Inter); Superficial (Super); Merkel Cells (Merk); Melanocytes (Mela); Langerhans Cells (LC); Migratory Dendritic Cells (MigDC); Mast Cells (Mast); also see Figure 1 legend. Illustration from (a) created with BioRender.com.

Next, we subclustered basal and SB JKs and SKs, finding that these distinct cell identities formed unique clusters using partition-based graph abstraction (PAGA; see Methods and Extended Data 6a-e). Pseudotime analysis confirmed terminal differentiation of SB JKs and SKs. We wondered how the gene expression pattern change might give insight into the differentiation and proinflammatory changes in SK/JKs. Along the differentiation trajectory in pseudotime, we observed altered differentiation patterns: *SPRR2A* and *SPRR3* decreased in JKs, yet in SKs, *SPRR3* and *SPINK7* decreased (Figure 3f and 3g). The largest difference occurred in cell signaling changes between JKs and SKs (Figure 3h and 3i). *CXCL1, CXCL8, IL1A, IL1B, CSF3, IL1RN*, and *CXCL3* all increased in JKs in periodontitis; alternatively, SKs were relatively less active. Overall, SK/JK stem cells and their progeny appeared to differently upregulate proinflam-matory, immunoregulatory, and innate immune chemotactic signatures at the expense of healthy cell differentiation patterns.

### Receptor-ligand analysis of epithelial and immune subpopulations reveals affected JK stem cells

Having confirmed cell identities of JKs and SKs and gained some insight into their differentiation and immune activity profiles, we further annotated innate and adaptive immune cell populations using CellTypist (see Methods and Extended Data 6f-g; Supplementary Table 1). We noticed that some DEGs in keratinocytes targeted both adaptive and innate immune cells. We became interested to understand their targets through recep-tor-ligand analysis. We utilized CellChat to understand cell-cell interactions via receptor-ligand interactions (see Methods), considering innate and adaptive populations separately (Figure 3j). In health, keratinocyte receptor-ligand interactions between the same and heterogeneous sub-populations were high, evidenced by larger “nodes”. This diminished in periodontitis. Immune cell communication patterns sharply increased, with larger nodes for multiple T and dendritic cell subpopulations (Figure 3j). We next investigated information flow from keratinocytes to innate and adaptive cells (Figure 3k). In health, both were generally dominated by periodontium structural support, including Laminin, Collagen, Junctional Adhesion Molecule (JAM), and Nectin signaling; in periodontitis, we observed increased cell signaling via CCL, EGF, PECAM1, and ICAM pathways, suggesting cell-cell communication shifts in disease, supporting a coordinated response.

With potential keratinocyte-adaptive and keratinocyte-innate axes identified, we next plotted predicted interactions between basal and SB JKs and SKs to understand cell-cell communication changes at the individual gene level for innate and adaptive populations (Figure 3l,m). In the innate plots, JK signaling to macrophages, neutrophils, and cDC1s dramatically increased; communication by GM KCs decreased. Basal cell signaling decreased; differentiated SB cell signaling generally increased. In the adaptive plots, basal SK/JK and GM KC receptor-ligand signaling also decreased, with most differentiated/SB signaling to natural killer T (NKT), tissue-resident memory T (TRM T), gamma delta T, and cytotoxic T cells. We quantified the confidence of receptor-ligand interaction between tooth-associated keratinocytes and immune cells (innate and adaptive, Extended Data 7a,b, respectively). *MIF*, associated with periodontal disease progression^37^, was a ligand (via *CXCR4* and *CD74*) in both the innate and adaptive panels: predicted interactions for MIF were strongest in JKs for neutrophils and macrophages (Extended Data 7a). We also found decreases in several innate and adaptive cell interactions between structural markers (Extended Data 7c,d; Supplementary Table 2). Overall, basal JK interactions were quite limited, suggesting these cells are especially affected in periodontitis; however, why these keratinocytes might be affected over nearby cells remained unclear.

### Polybacterial coinfection of human keratinocytes is diverse but frequent in periodontitis

Considering polymicrobial dysbiosis drives periodontitis in susceptible hosts6, we hypothesized these primarily SK/JK phenotypes partially arose from spatially-focused periodontal pathogen (“periopathogens”) infection profiles. We first used a *16S* rRNA ISH probe common to all bacteria^38^ in healthy and diseased tissues (Figure 4a). In health, *16S* was frequently detected in suprabasal keratinocytes; however, in periodontitis, we noted higher counts of *16S+* basal keratinocytes and stromal cells, with generally higher burdens in JKs. We wondered if we could reveal cell-specific, species-specific bacterial burden by adapting and tailoring the “Single-cell Analysis of Host-Microbiome Interactions” (SAHMI; see Methods) algorithm to identify bacterial reads from single-cell datasets (Figure 4b; Supplementary Table 2). We first looked at healthy samples, identifying few cell-microbial associations (Figure 4c); yet, in diseased samples, we observed many more bacterial reads per cell. Well-known periopathogen *Porphyromonas gingivalis*^39^ had the largest increase–nearly 200-fold–in keratinocytes and even larger increases in other immune cell populations (Figure 4d,e). Other well-known periopathogens *Leptotrichia sp.*^40^, *Treponema (T.) denticola, T. medium, T. vincentii*^41^, and recently associated periopathogen *Pseudomonas aeruginosa*^42^ had >2-fold increases. These data suggested cell-specific enrichment for oral microbes in disease.

**Figure 4.**
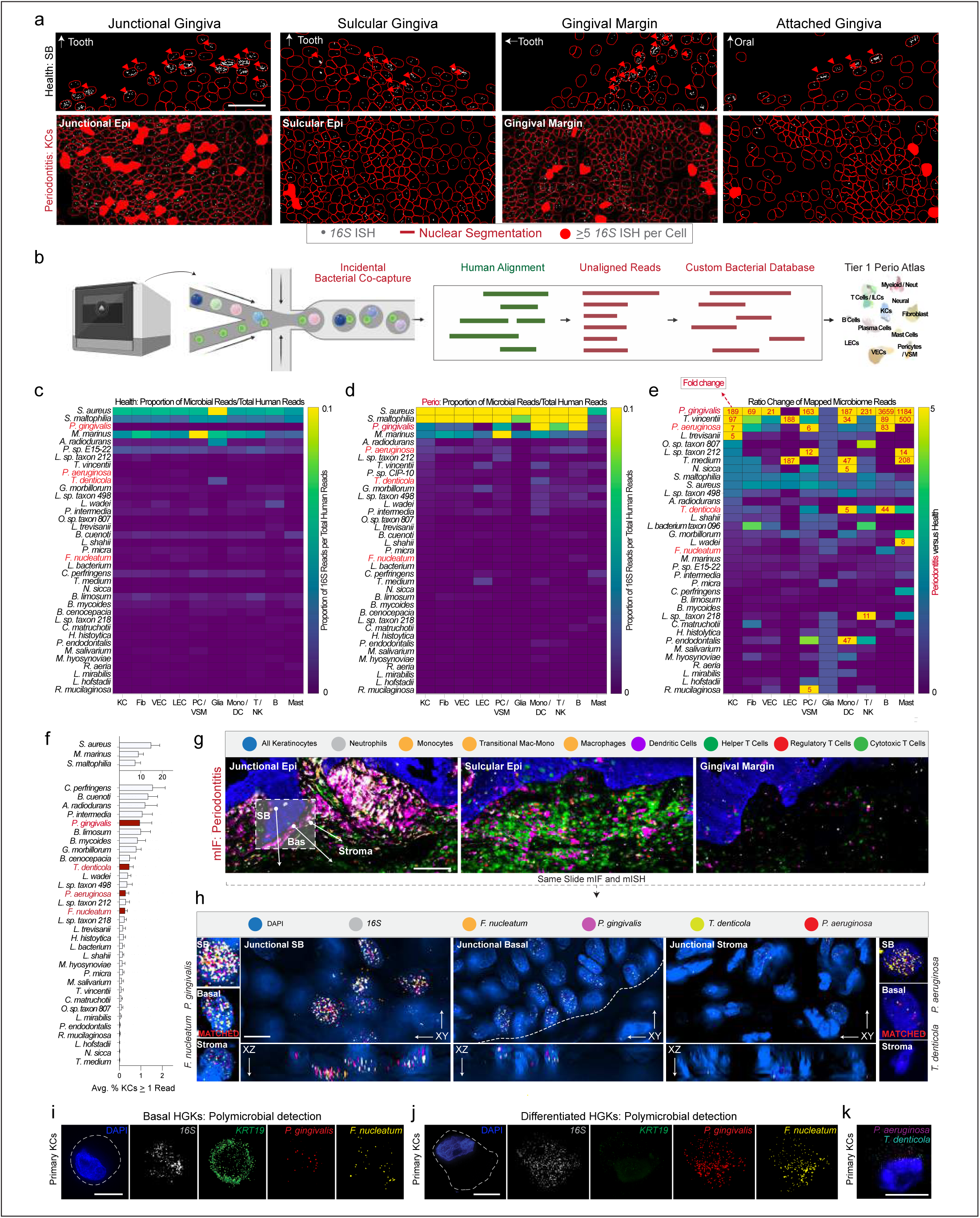
Discovery of polybacterial coinfection of human keratinocytes in vivo and in vitro. (a) Using a 16S in situ probe, we segmented cells using StarDist in health and disease tissues. We found bacterial signals primarily focused on the most terminally differentiated suprabasal keratinocytes across each region of the oral and tooth-associated keratinocytes. The most bacterial burden was found to be in JKs. In disease states, epithelial barrier integrity appears compromised as we see more stromal and epithelial stem cells associated with bacteria, especially in JK and SKs. (b-f) (b) Using a modified pipeline for single-cell analysis of host-microbiome interactions (SAHMI) and a custom Kraken2 database, we used the unmapped reads from our integrated single-cell periodontitis meta-atlas. (c) Using a broad Tier 1 annotation of cell types, this analysis revealed 37 distinct species across Keratinocytes (KC), Fibroblasts (Fib), Vascular Endothelial Cells (VEC), Lymphatic Endothelia Cells (LEC), Pericyte/Vascular Smooth Muscle (PC/VSM), Glial Cells (Glia), Monocyte/ Dendritic Cell Lineages (Mono/DC), T and Natural Killer Cells (T/NK), B Cells (B), and Mast Cells (Mast). In health, using a fraction of microbial reads to the averaged reads per human cell class, we find low read counts across most of the 37 bacterial species. (d) In disease cells, we find large shifts in associated reads. (e) Performing an analysis of c,d, we find dramatic increases in many bacteria comparing disease to health, with large increases in known periopathogens (i.e., *P. gingivalis, T. vincentii, P. aeruginosa [P. sp. CIP-10]*). (f) Focusing on all keratinocytes, we find variable numbers of bacteria associated with these cell types, ranging from 0.1% to 15% of all KCs. (g-h) (g) Utilizing broad cell classification of our multiplex immunofluorescence data (mIF; Figure 2), we show the innate versus adaptive immune foci in disease. (h) By using the same mIF slides and targets predicted from our SAHMI pipeline, we apply in situ hybridization against *16S* and four periopathogens (*F. nucleatum, P. aeruginosa, T. denticola,* and *P. gingivalis*) and find polybacterial coinfection of all four species in some epithelial stem cells of the JKs using Nyquist optimized, three-dimensional imaging. (i-j) (i) Using primary human gingival keratinocytes, we find polybacterial coinfection in basal (i) and suprabasal KCs (j) for *F. nucleatum* and *P. gingivalis*. We also find this phenomenon for *P. aeruginosa* and *T. denticola* (k). Abbreviations: see Figure 1 legend. Scale bars: 100 μm; 50 μm; 10 μm; 5 μm. Illustration from (b) created with BioRender.com.

Because some periopathogens—*P. gingivalis, F. nucleatum*, and *T. den-ticola*—have been found intracellularly in keratinocytes, fibroblasts, and endothelial cells^43^, we hypothesized that SK/JKs may harbor these microbes intracellularly. Using healthy and diseased keratinocytes, we first looked at the average number of cells harboring at least one bacterial read per barcode, finding enrichment of key periopathogens in 0.5-2% of all barcodes (Figure 4f). We then created an ISH panel against 4 of the detected periopathogens, with the *16S* rRNA probe as a control. We re-used the PhenoCycler-Fusion tissues and re-probed for microbial targets (Figure 4g,h) revealing intracellular pathogenesis of all 4 periopathogens in 3D—occasionally all four in the same JK epithelial stem cells. While 500 species have been detected at this critical niche in humans^44^, obligate anaerobes like *T. denticola* localized to the deepest part of JK at the tooth-soft tissue interface; facultative anaerobes like *P. aeruginosa* could be found in JK and SK cells—but in few cells elsewhere (Extended Data 8a).

This polybacterial intracellular coinfection phenotype was surprising and provided the first known evidence of intracellular residency of at least 4 pathogens within the same cells. To show polybacterial coinfection extended beyond a gingival phenotype, we similarly profiled human tonsils, finding *P. gingivalis* and *F. nucleatum* in tonsil basal keratinocytes (Extended Data 8b). Further, we used HGKs and found evidence for polybacterial coinfection at passage 2 (P2) through P4 of the same periopathogens (Figure 4i), finding the same phenomenon in differentiated cells, which are much larger than basal stem/progenitors (25-50 versus 5-10 µm, respectively). We consistently observed proportional *16S* count increases with cell cytoplasm size, suggesting actively replicating bacterial species within these keratinocytes (Figure 4j). We also found evidence of the same polybacterial coinfection for *P. aeruginosa* and *T. denticola* (Figure 4k). By linking single-cell, multiplex (m)IHC, mISH, and primary cells with cell-specific markers, we predict that polybacterial intracellular coinfection is likely more common than previously appreciated.

### Polybacterial coinfection drives sustained proinflammatory phenotypes in vitro and in vivo

We suspected that the SK and JK phenotypes we observed in the sin-gle-cell data could partly be attributed to polybacterial coinfection. Because we consistently saw cell-specific signatures, we termed this epithelial cell signaling response as “epi-kines/keratokines” i.e., cytokine upregulation in response to challenge. To test this in vitro, we cultured HGKs that displayed varying levels of polybacterial coinfection and challenged them with 20 µg/mL of *P. gingivalis* LPS for 48h, then fixed and immobilized them for analysis using another custom ISH panel against 9 keratokines, combining 3 per cycle with 16S (Figure 5a). We hypothesized that polybacterial coinfection alone could induce sustained keratokine profiles, and some signatures would synergize with extracellular LPS.

**Figure 5.**
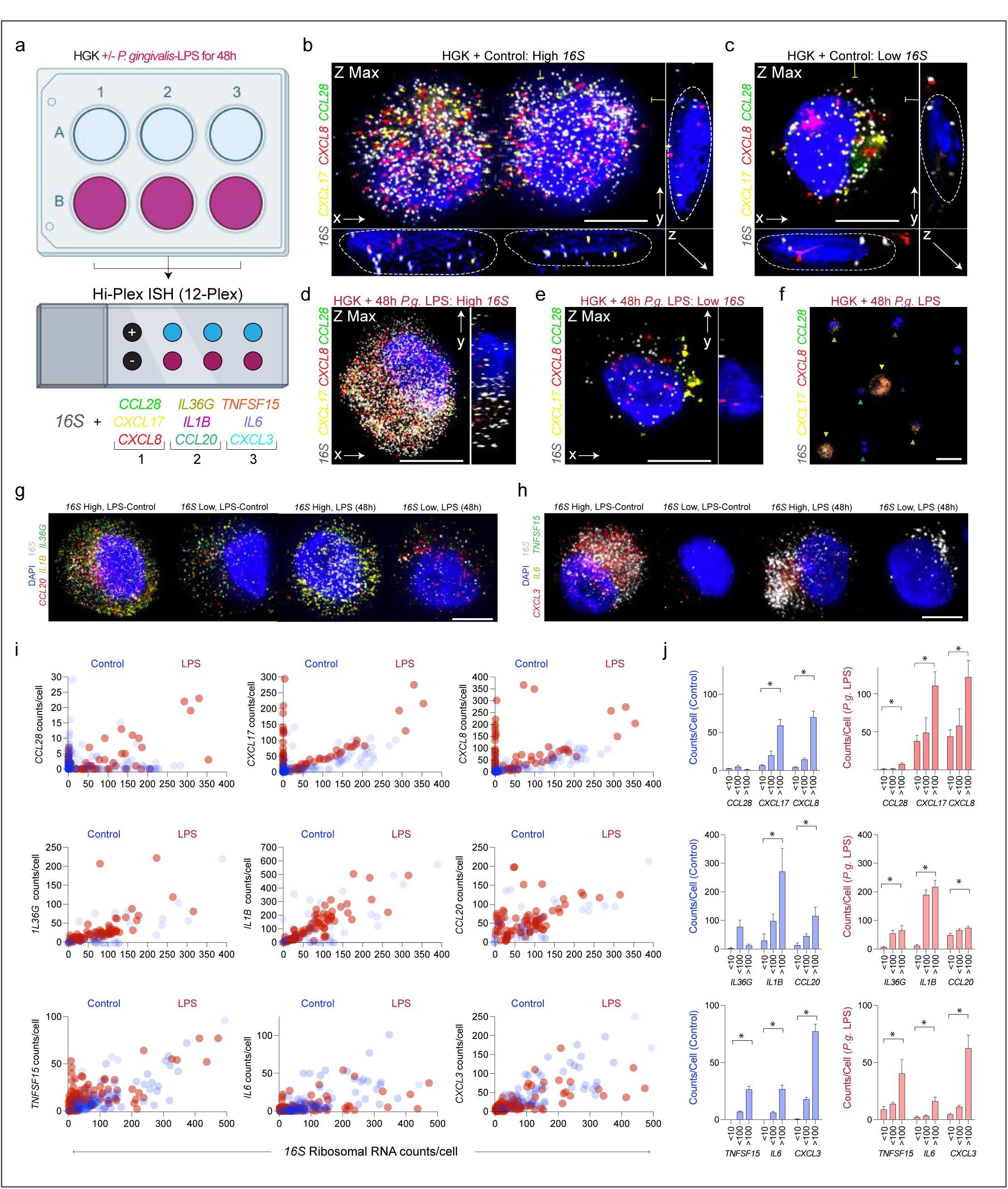
Polybacterial coinfection uniquely affects human gingival keratinocytes and synergizes with LPS. (a) Human gingival keratinocytes (HGK) harboring polybacterial coinfection were cultured with and without LPS (P. gingivalis). These cells were assessed for *16S*, and cytokines were predicted to increase in disease states using Hi-Plex in situ hybridization in three rounds. (b-h) (b) HGKs were imaged in xyz using Nyquist-optimized parameters and showed a correlation of *16S* burden and cytokine mRNA production. This happens without LPS (b-c) and increases in HGKs with LPS. The first round of images highlights this phenomenon for *CCL28, CXCL17,* and *CXCL8*. (g-h) (g) Examples of *16S* and the other cytokine panels (*IL36G, IL1B, CCL20*; (g)) and (*TNFSF15, IL6, CXCL3*; (h)) with and without LPS. (i-j) (i) Quantification of all cytokines considering *16S* signals per cell. For *CXCL17, CXCL8, IL36G, IL1B, CCL20, TNFSF15, IL6,* and *CXCL8*, polybacterial coinfection elicits mRNA expression without LPS. Only *CCL28*, which is known to be chemoattractant to T and B cells, is downregulated with increasing bacterial burden. Binning cells in *16S* groups with and without LPS challenge, *CXCL17, TNFSF15,* and *CXCL8* expression demonstrates a synergy between LPS and polybacterial co-infection. Abbreviations: In situ hybridization (ISH); Human Gingival Keratinocytes (HGK); Lipopolysaccharide (LPS). Scale bars: (f) 100 μm; 10 μm. p<0.05, paired Student’s T-test (j).

ISH showed that in both LPS challenged and unchallenged HGKs, the cells maintained a resident bacterial microenvironment, evidenced by *16S* signal (Figure 5b-h). In unchallenged cells, *CXCL17* and *CXCL8* expression numbered in the tens per cell, but in the hundreds in challenged cells. Further, we unexpectedly found a direct linear correlation between the number of *16S* and *CXCL17* and *CXCL8* counts per cell—not found for *CCL28*, though LPS induced more higher-burden cell expression that synergized linearly with *16S* counts. *16S* burden alone accounted for increased *IL36G, IL1B, CCL20, TNFSF15,* and *IL6* (Figure 5i,j). Overall, polybacterial coinfection drove single-cell phenotypes. These cells, however, were harvested from an otherwise healthy donor, so we sought to understand this phenomenon in vivo.

To investigate single-cell, polybacterial coinfection in health and periodontitis in situ, we added two additional keratokines to the panel and performed three consecutive imaging rounds, aligning the images to simultaneously assess all 12 probes at single-cell and spatial resolution (Figure 6a). We segmented stroma and basal/SB keratinocytes at the four ROIs (Figure 1,2), subdividing each tissue into 12 segments. We generated custom scripts to run 12-plex ISH analyses from these ROIs, qualitatively noting *16S* signal correlated with strong keratokine upregulation, irrespective of the ROI (Extended Data 8c). We quantified this and performed statistical analysis. *CXCL8, CXCL17,* and *CCL28* all increased with increasing *16S* (Extended Data 8d-f). Simultaneous keratokine comparison revealed highly infected stromal cells and keratinocytes displayed cell-specific phenotypes (Figure 8b). Other markers were enriched in suprabasal cells or the stroma specifically. *IL1A, IL1B, CXCL1, CXCL3,* and *IL6* displayed some correlation with highly infected cells (Figure 8c)—mostly suprabasal cells, likely explaining pseudotime plot differences in periodontitis (Figure 3).

**Figure 6.**
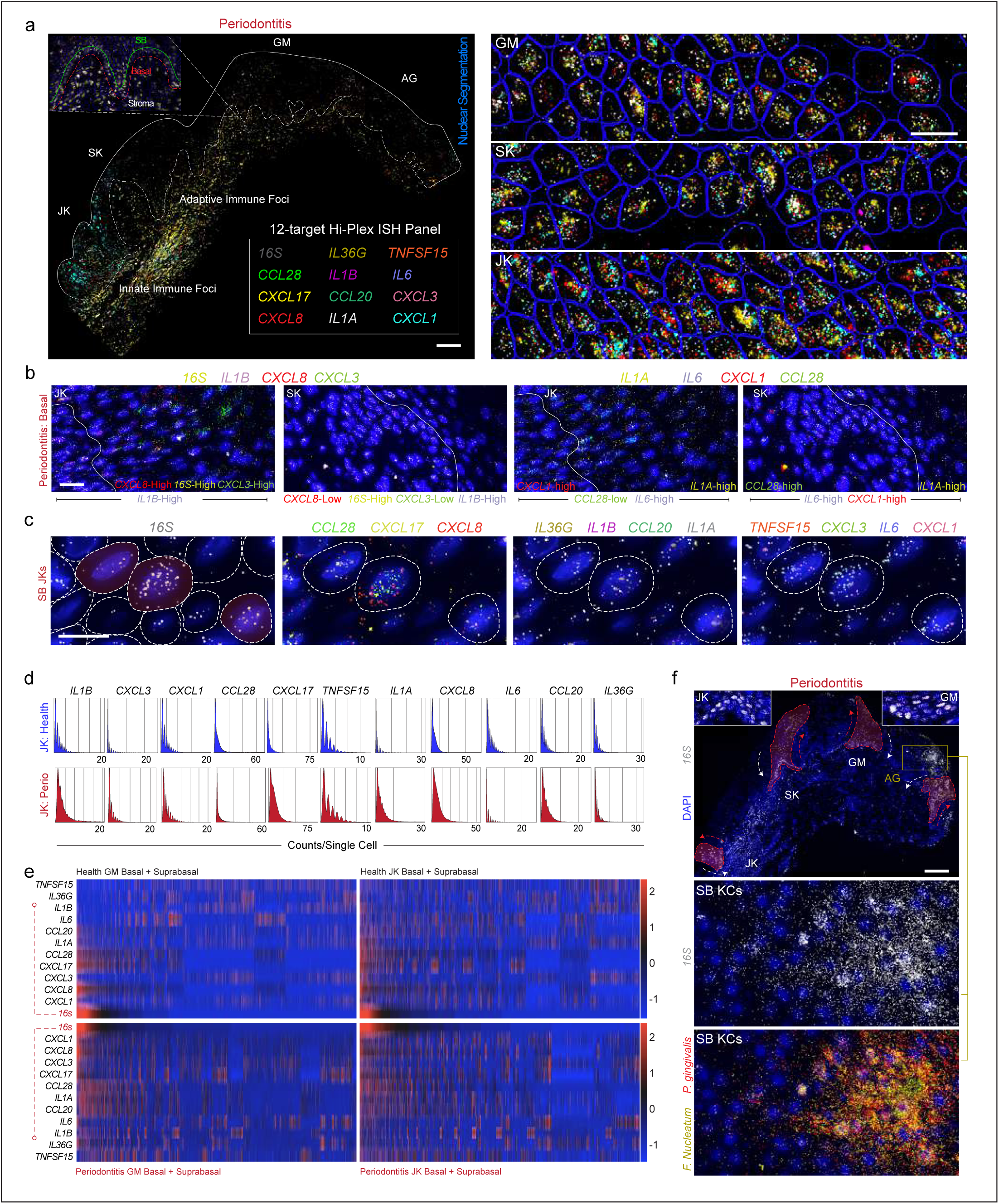
Polybacterial coinfection of human keratinocytes in vivo similarly affects cells in health and disease. (a) 12-plex in situ hybridization panel of mRNA targets and 16S were overlaid using Warpy. All 11 cytokines are shown simultaneously in GM, SK, and JKs. (b) Without *16S*, there is a distinct patterning of each cytokine. Some, such as *IL1B,* are broadly expressed in epithelial and stroma. Others such as *CXCL8* and *CXCL3* appeared to be cell-specific and enriched in JK over SK. (c) Considering infected keratinocytes at a cell-specific level, we find that *16S* alone is positively associated with most cytokines in health and disease states. (d) There appears to be polybacterial infection patterns in disease that spread to all epithelial regions, including in terminally differentiated keratinocytes of the attached gingiva (i.e., keratinized mucosa). Epithelial stem cell infection was found in each region. (e-f) (e) Ridge plots (n=3) from JKs in health and disease were quantified without consideration of 16S, showing general increases in *IL1A, IL1B, CXCL8, CXCL17,* and *CCL20*. Assessing both JKs and GM, all keratinocytes were plotted on a normalized heatmap relative to 16S expression, and we quantify that *CXCL8, CXCL17, CCL28, IL1A, IL1B*, and *CCL20* are associated with microbial burden in healthy GM keratinocytes. In JKs, nearly all cytokines are positively associated with microbial burden in heath, suggesting that some bacteria (i.e., gram-positive in GM, gram-negative in JK) may have cell-specific effects in vivo. (f) Patterns in disease between GM and JK align in disease, suggesting an effect by periopathogens as shown in D. Scale bars: (a,d) 100 μm (insets; 50 μm); (b-c) 25 μm. Abbreviations: In situ hybridization (ISH); Human Gingival Keratinocytes (HGK); Lipopolysaccharide (LPS); also see Figure 1 legend. Illustration from (a) created with BioRender.com.

Broadly, when comparing health and periodontitis, we found upregulation of keratokines by higher RNA transcript numbers per cell (Figure 8d). IL1 superfamily members *IL1A, IL1B,* and *IL36G* were upregulated in periodontitis in both the basal and suprabasal layers of the JK; further, though previously reported, we observed sole concomitant expression of these genes in the peri-junctional stroma. These observations held in *CXCL1* and *CXCL3* expression—both innate immune cell chemokines. We reordered keratokine expression heatmaps by *16S* burden in the JK and distant GM regions (Figure 6e). 16S-high burden directly correlated with high *CXCL1, CXCL3, CXCL8,* and *IL1A*, with *IL6* additionally corresponding to basal and suprabasal layer invasion—in either ROI, suggesting though JKs may harbor periopathogens more often due to their specific niche, other cells can acquire this phenotype. This was more pronounced for CC and CXC motif chemokines in either region.

In disease states, we further evaluated the epithelial stem cell and differentiated progeny affected by this polybacterial coinfection phenomenon, revealing pseudo-lineage tracing patterns in AG, GM, JK, and SK regions (Figure 8f), reminiscent of lineage patterns in mouse oral epithelial stem/ progenitor cells^30,45,46^. This is important because these cell types are persistent in the basal layer, giving rise to differentiated progeny for many cell divisions, potentially passing on these bacteria to subsequent stem and suprabasal cell generations. Furthermore, the current treatments for periodontal disease, such as scaling/root planing, periodontal surgery, and systemic antibiotics are unlikely to clear these pathogens, setting up a lifespan-sustained persistent infection. Considering periopathogen increases in biofluids like blood (i.e., bacteremia^47^) and saliva^48^ as well as epithelial stem/progenitor longevity in the basal layer^49^, this could be a body-wide phenomena associated with inflammaging.

## DISCUSSION

The promise of precision medicine is for earlier disease intervention, improved clinical outcomes, and a general improvement in quality of life through extending both the life-and healthspan^50^. To date, periodontitis has not benefited from this promise for several reasons related to its complex host genetics^51^, polymicrobial heterogeneity^52^ along an anteroposterior axis caused by swallowing^53^, systemic periodontium effect from other chronic/genetic diseases^54^, and even same-patient differences between “asymmetric bursts” and linear disease activity^55^. Our study highlights challenges to implementing precision aproaches in periodontal medicine may partially arise from an incomplete pathophysiological complexity understanding at a single-cell and spatial biological level^56^. Even with large-scale efforts i.e., the Tabula Sapiens project (∼500k cell; 24 body-sites)^57^, more work remains to be done identifying and validating new cell types—here, in human oral cavity microniches, specifically in identifying niche-resident cells and their impact in concert with other structural cell types^21^. Furthermore, spatial biology is in its ascendency, helping to determine new cell identities and states^58^ and pinpoint cell locations and functions in health and disease^21^. Since this is one of the first studies to use spatial ‘omics assays in the oral tissues, significantly more work remains to understand each niche in more detail to support future precision initiatives.

Additionally, despite a singular focus on one specific niche within human tissues, much remains to be understood in adult homeostasis, aging, and in development prenatally or during childhood. This study highlighted likely as-of-yet undiscovered cells in periodontitis. Some rare^59^ or difficult-to-sequence cell types like eosinophils^60^ are underrepresented in these atlases, making validation difficult until enrichment for sequencing or targeting through in situ approaches^61^. In our atlas, tooth-associated keratinocytes (SK/JK) represent a rare epithelial cell population (1.1%; stem/progenitor and SB fractions much lower); however, even rarer are epithelial-resident *KRT20+; ATOH1+* Merkel cells (0.02%) and *MLANA+* Melanocytes (0.1%)—not found in high enough proportions even when combining four studies. Basophils, eosinophils, innate lymphoid cells, and mineralized tissues (osteoblasts, osteoclasts, osteocytes, cementocytes, etc.) were not annotated here; peripheral nervous system contributions via myelinating or non-myelinating Schwann cells were also undersampled (0.05%). Thus, we refer to this as a “draft v1” of the atlas; with more datasets and multimodal sampling of this niche, more will be refined, annotated, and learned that can benefit precision approaches through digital “tooth” modeling of druggable targets at a single-cell level via available therapeutics (i.e., heart; drug2cell^62^). Adapting this in a spatial context will also be necessary to overcome these described challenges for precision approaches for periodontal diseases.

This original study shows the “spatial” nature of periodontal disease, due to tissue orientation challenges and microniche breakdown in disease states. Here, we link historical knowledge and approaches to annotate new keratinocytes and describe their function. This study singularly focuses on keratinocytes; more work will be forthcoming about other structural immune influences. As a first focus, we knew that the oral/gingival epithelium is comprised of *K14*/KRT14-high epithelial cells, yet some KCs are also *K19*/KRT19-high in the gingival “pocket”^11^. Previous studies showed common cell ontology class representation in each scRNAseq dataset; using Cellenics® enabled the integration and collaborative, harmonized annotation of these datasets as well as KC discovery and validation. It will be important to design future studies with both Tier 1 and Tier 2 clinical metadata, including detailed descriptions of sample origins (CCF29) for dataset harmonization to allow for host impact discovery of this chronic disease.

Our study highlights another challenge to precision periodontal medicine with body-wide implications in many diseases. After cell annotation and spatial validation, our multiomic toolkits gave us insight into cellular programming shifts in single-cell polybacterial intracellular coinfection phenotypes only when combined. Utilizing scRNAseq, (m)IHC, (m)ISH, and cell culture, we linked altered differentiation patterns and upregulated keratokines in these new cells to a specific host-microbe-interaction cell state. JKs demonstrated biased signaling for macrophages and neutrophils (*CXCL1/3/8*), whereas SKs biased signaling toward T/ NK and B-cells (*CCL20/28*), correlating with polybacterial intracellular coinfection, likely with structural immunity correlations with peri-junctional and sulcular stromal foci and peri-vascular microniches, which warrants further investigation. Further, while some studies suggest tumor-specific microbiomes^64^, these relationships in our study could only be discerned at a cell-specific and niche-specific level. While polymicrobial coinfections are not thought to occur with multiple viruses at a single-cell level (i.e., superinfection exclusion^65^), polymicrobial (bacterial-bacterial; bacterial-viral) coinfection phenomena cause multiple inflammatory diseases across the body^66^. Recent work has implicated numerous bacteria such as periopathogen F. nucleatum in colorectal and oral cancers38 with similar immune invasion^67^ found here; similar to our periodontitis findings, this cancer study also found *Porphyromonas, Streptococcus,* and *Leptotrichia* genus in colorectal cancers, further supporting and oral-systemic “intermucosal” linkage^44,68^.

We termed this observed phenomenon “polybacterial” intracellular coinfection because we observed and analyzed these phenotypes at a single-cell level in vitro and single-cell and spatial level in vivo with 3D imaging. While many diseases present with polymicrobial infections generally, this has been historically described at a tissue level across bacteria, viruses, fungi, and parasites69. While P. gingivalis had broad cell tropism in disease using single-cell metagenomics (Figure 4), Treponema sp. appeared to have lymphatic endothelial cells (LEC) tropism, with other enrichment patterns including P. endodontalis in pericytes/vascular smooth muscle cells (PC/VCM) and monocytes/dendritic cells and R. mucilaginosa in PC/VSM. Periodontal disease cell-specific bacterial infection validation was focused on keratinocytes, with minimal stromal population exploration here. Our single-cell metagenomics approach has generated more hypotheses than approached, especially bacteria-driven immunosuppressed phenotypes in disease at a single-cell and spatial context, likely differing across age, sex, and genetic ancestry. Our findings of bacterial infection in both tissue and primary cell lines further ask new questions about current long-term periodontitis treatment effectiveness70. Finding new ways to clear intracellular pathogen residency will allow true restoration of the tissue niche.

## Supporting information

Supplemental Table 1: Clinical metadata, Tier marker genes, and Gene expression change.

Supplemental Table 2: CellphoneDB, CellChat, and Bacterial enrichment.

---

**Extended Data 1.**
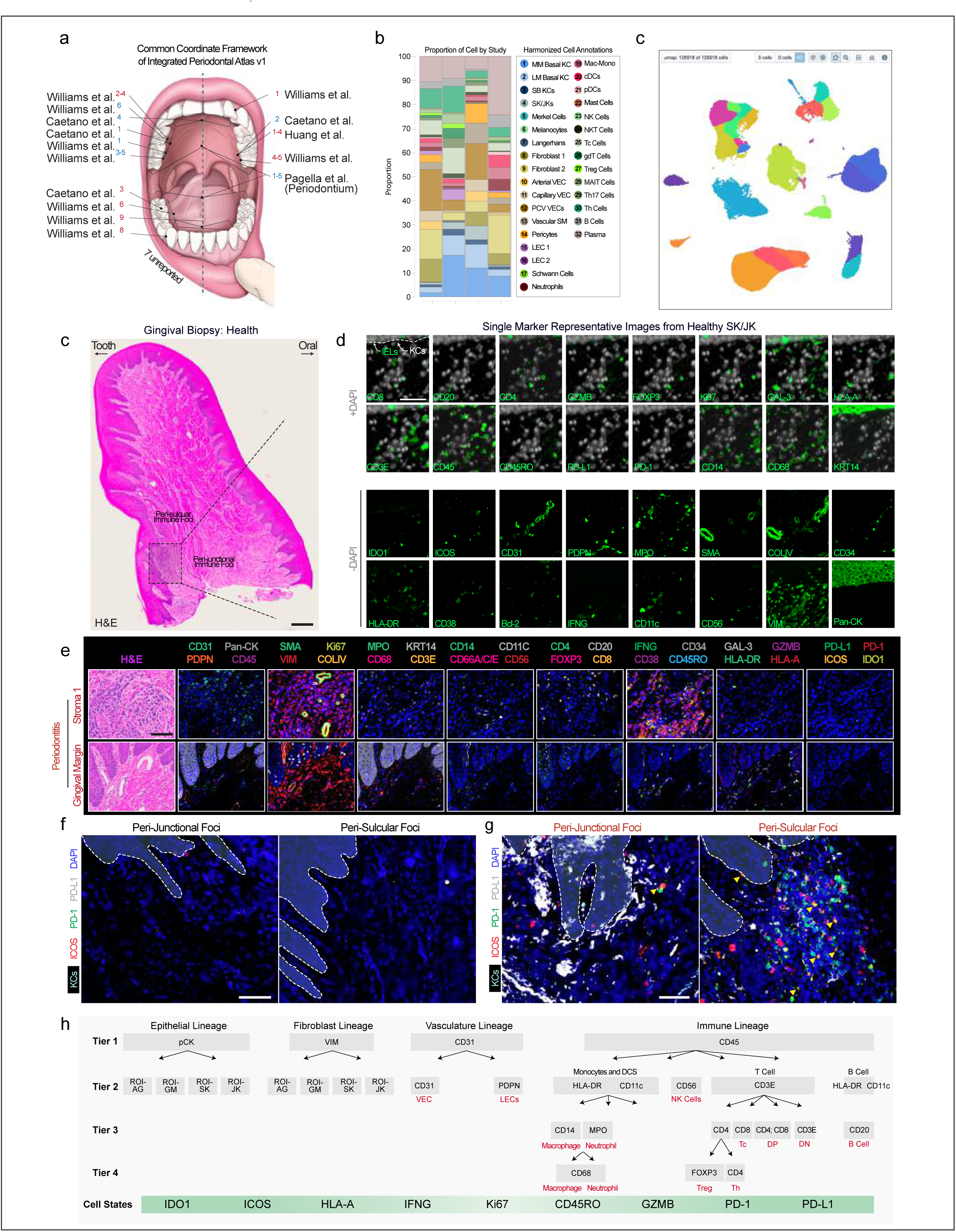
Metadata and Spatial Analyses of the Integrated Periodontitis Meta-atlas. (a) Common coordinate framework annotations were included from each study here visually and in Supplementary Table 1. Some harvest sites were unknown (∼20%), and others were taken from more broad regional characterizations than a single tooth (i.e., anterior or posterior; ∼25%). To be in alignment with the Human Cell Atlas and future integration efforts, standardized Tier 1 and Tier 2 metadata will be required to support future drafts of this atlas (see Supplementary Table 1 for template). (b) As referenced in Figure 1, no single study contained all the cell types annotated in this meta-atlas; however, comparing the studies using the harmonized cell annotations, these four studies are not grossly different when comparing relative cell proportions. (c) For this study, Cellenics®, which is an open-source tool for single-cell RNA sequencing analyses (https://github.com/hms-dbmi-cellenics), was used for integration, data analysis, and some plot generation. For public use, Cellenics® and cellxgene were linked, conserving the UMAP coordinates in the cellxgene space. Metadata from Supplementary Table 1 was incorporated into cellxgene (https://cellxgene.cziscience.com/) to further enhance public utility. (c) Orientation of health tissue with tooth and oral facing sides allows for peri-junctional immune foci characterization also in health. (d) Validation of single markers in the health periodontal niche. As expected, there are few adaptive immune cells and minimal exhaustion phenotypes expressed in healthy tissues, though the peri-junctional space does express some of those identities and states in health. (e) Additional zoomed-in areas around the periodontal lesion from Figure 2c further validate expression patterns of antibodies in areas that do not show immune cell infiltration from H&E sequential sections. (f,g) Examples of the healthy (f) and diseased peri-junctional (g) and peri-sulcular immune exhaustion phenotypes quantified in single marker analysis (Figure 2f, g) and spatial immune foci analysis (Figure 2l-n). (h) Tier assignment algorithm for multiparameter cell type assignment and cell state analysis used n Figure 2. Abbreviations: see Figure 1 legend. Scale bars: 250 μm; (b-d) 50 μm.

**Extended Data 2.**
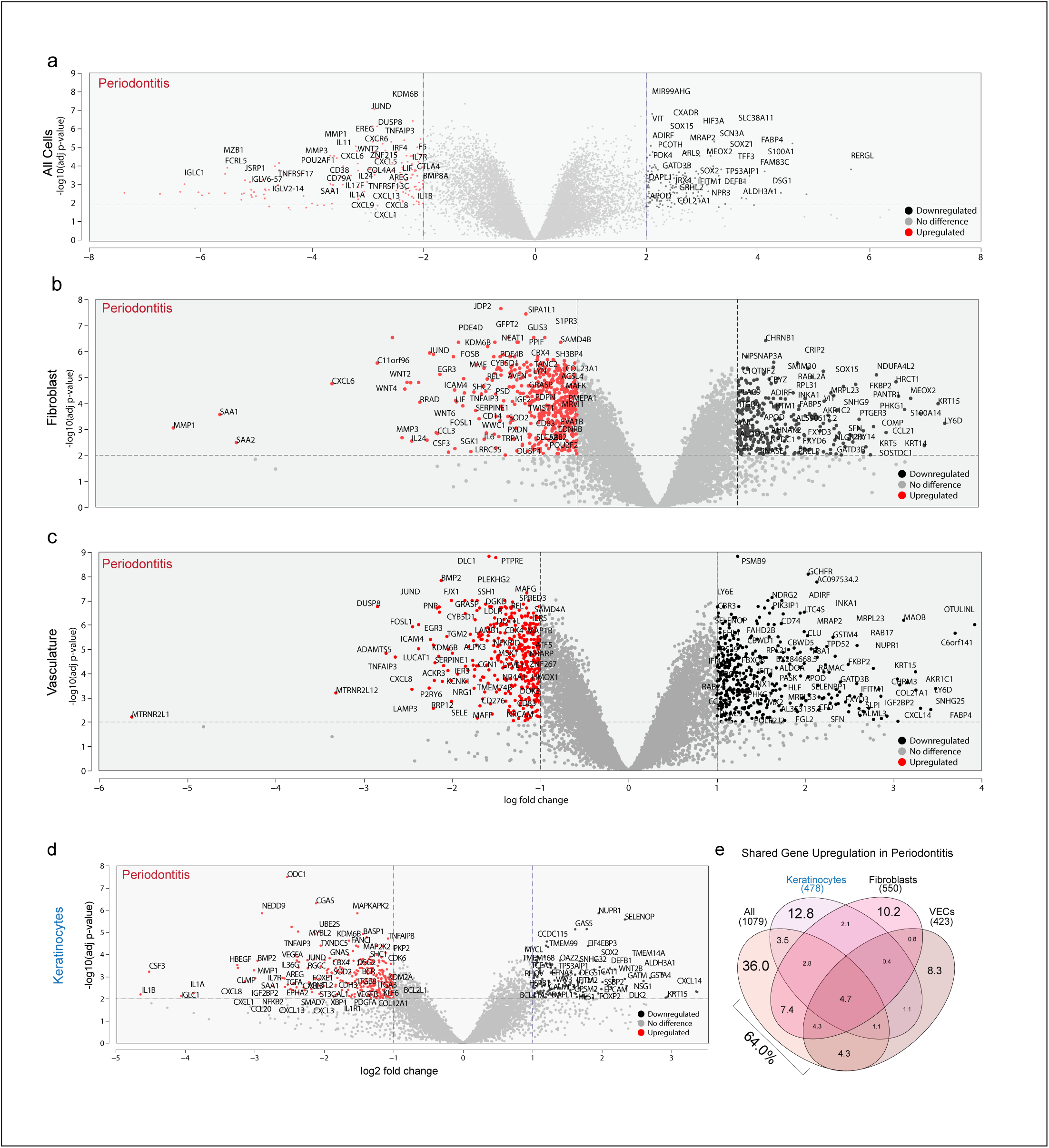
Investigation of the “Structural Immune” Contribution to Periodontitis using Pseudobulk and Cell-specific Analyses. (a-d) Pseudobulk analysis of differentially expressed genes (DEGs) in periodontitis using all (a) and Tier 1 cell annotations of (b) Fibroblasts (c) Endothelial Cells/Vasculature, and Keratinocytes (d) are shown via volcano plots. Only some DEGs are highlighted; the full list is in Supplementary Table 1. (e) Using a Venn diagram revealed that many genes in these three cell types appear in the pseudobulk DEGs from (a). Abbreviations: see Figure 1 legend. Illustration from (b) created using Interactivenn.net.

**Extended Data 3.**
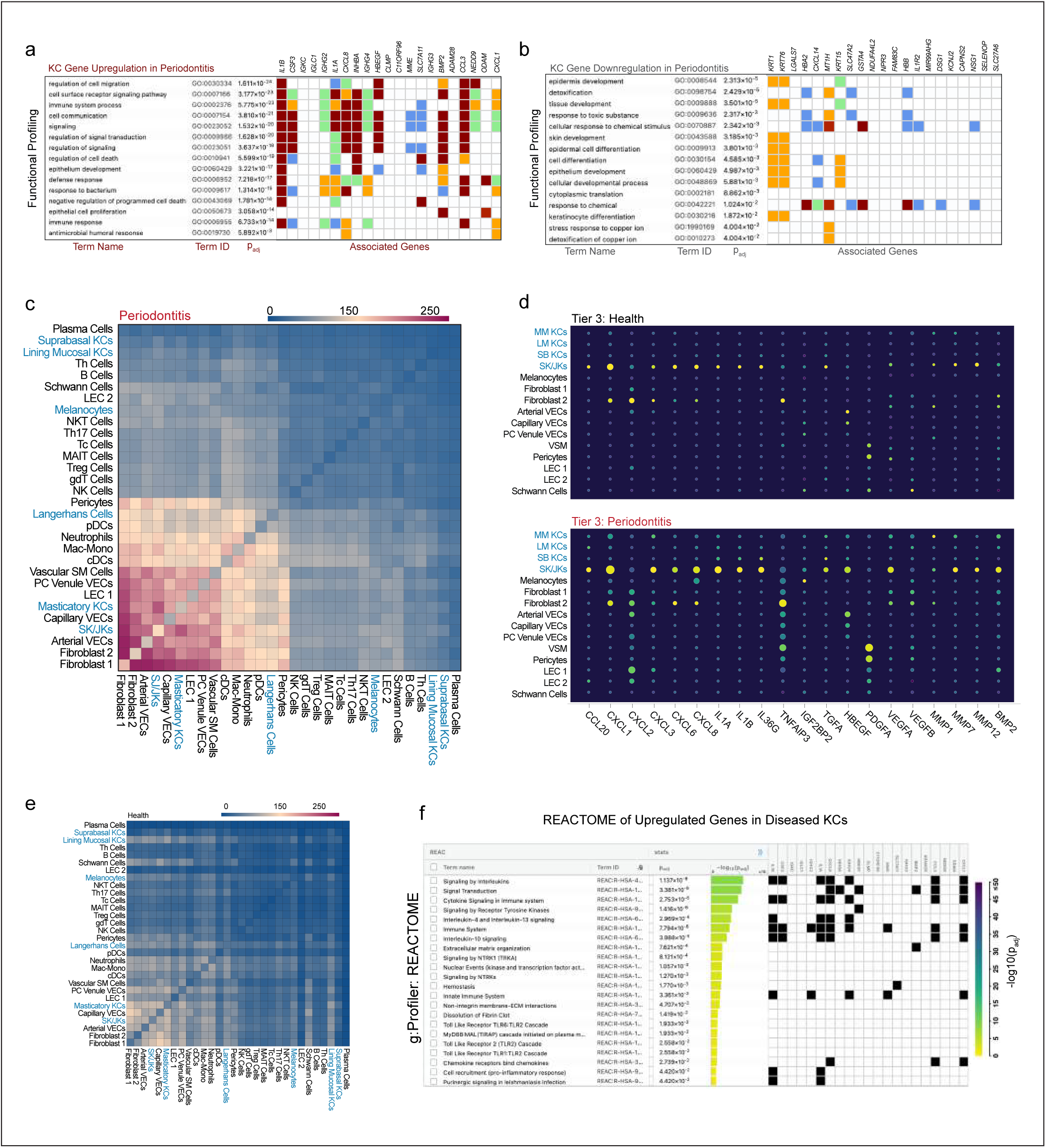
Keratinocytes play a prominent role in the host response to periodontitis. (a-b) (a) Assessing biological pathways using g:Prolifer, key processes that are upregulated in disease include cell migration, cell signaling, cell death, and cell responses to bacteria. (b). Key processes that are downregulated include tissue differentiation and development, protein translation, and stress responses. This is just an example of the g:Profiler data and key genes attributed to these pathways; however, these data suggest an active immune signaling role for keratinocytes in periodontitis besides just wound healing. (c) Using CellPhoneDB and the integrated periodontitis meta-atlas, all Tier 3 cell types were assessed for inferred receptor-ligand interactions. As expected, the most active cell types in periodontitis predicted include fibroblasts, vascular endothelial cells, and keratinocytes which are also predicted to interact with innate immune cells over adaptive immune cell populations. This full list is included in Supplementary Data 2. (d) Using dot plots to assess some key signaling genes in periodontitis as shown by the DEGs from (Extended Data 2), SK and JKs demonstrate higher expression of these genes, even in health. Furthermore, the patterns appear complementary to vascular and fibroblast populations, suggesting these cells may be responding to something in the clinical “healthy” state. (e) CellPhoneDB on the same scale as (c) shows less receptor-ligand activity in health but SK and JKs are relatively higher and focused on interactions between vascular endothelial cells (VECs), lymphatic endothelial cells, and Schwann cells— and less on fibroblasts and other cell types. This full list is included in Supplementary Data 2. (f) Using g:Profiler, the reactome of keratinocytes further emphasizes their active role via cytokine signaling, immunoregulation, and immune cell recruitment which was observed as presented in Figure 2. Abbreviations: see Figure 1 legend.

**Extended Data 4.**
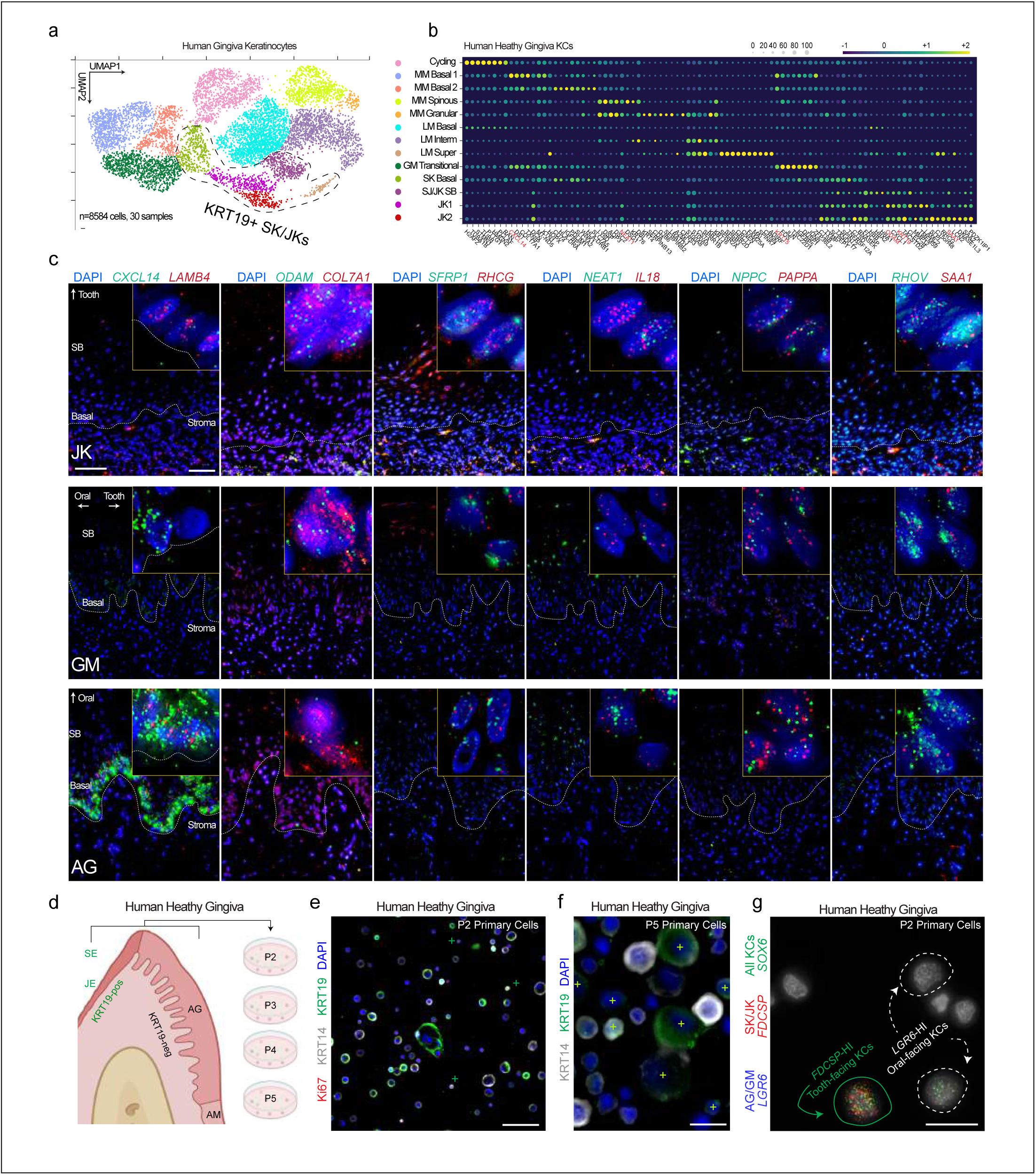
Discovery and validation of novel human keratinocyte populations in vivo and in vitro. (a-b) (a) Using Cellenics®, all keratinocytes (*KRT14+*) were subclustered from the integrated periodontitis meta-atlas (30 samples, 8584 cells) and assigned annotations (b) based on Louvain clustering. (b) Cell signatures for these populations are plotted and included in Supplementary Table 1. (c) Using these signatures, a HiPlex panel of 12 markers was used to profile all keratinocytes from this niche (AG, GM, and JK here; SK and AM as in Figure 1). Markers like *CXCL14* mark the AG basal epithelium in the opposite pattern of *KRT19, ODAM, RHCG, IL18,* and *SAA1*. Furthermore, keratinized epithelial populations (AG) share expression of *CXCL14, KRT15,* and *LGR6, SFRP1.* (d) Primary human gingival keratinocytes were cultured over multiple passages and (e) *KRT19*-high basal and larger suprabasal keratinocytes are found mixed population at (e) first passage and over (f) multiple passages. (g) Using RNA ISH and additional markers, cell subpopulations can be identified, suggesting a heterogeneous model of tooth-facing and oral-facing keratinocytes can be utilized for assays and that these markers are more likely cell identities than cell states (see Figure 5). Abbreviations: Passage (P2); see Figure 1 legend. Scale bars: (c, e) 50 μm; (f) 25 μm; (g) 10 μm. Illustration from (d) created with BioRender.com.

**Extended Data 5.**
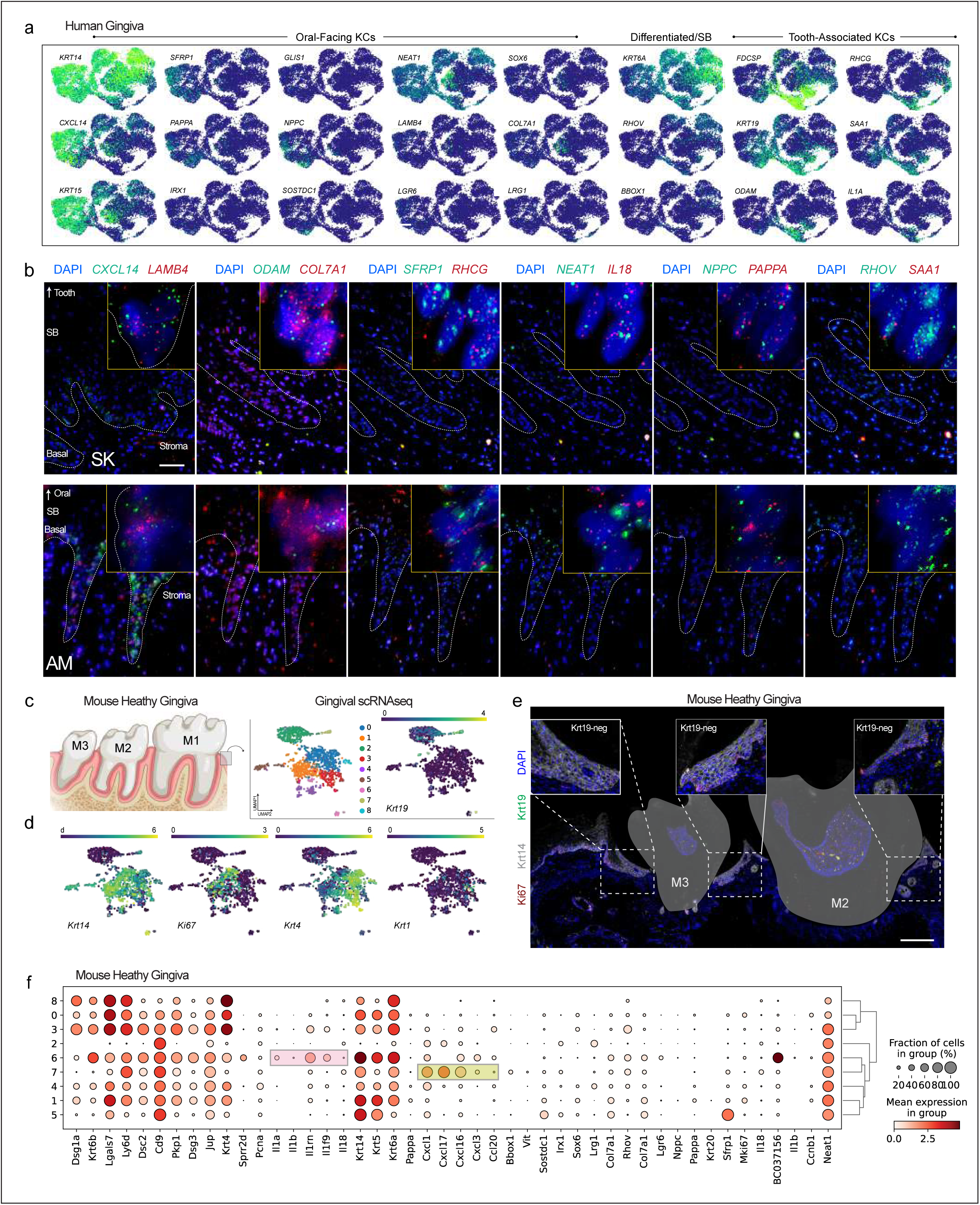
Human gingival keratinocytes contain distinct subpopulations. (a) Using UMAPs to demonstrate cell marker enrichment, keratinocytes are defined by *KRT14* expression. JK and SKs are defined by *KRT19, FDCSP, RHCG, SAA1, IL1A,* and *ODAM.* (b) Validation of markers using a 12-plex ISG in non-keratinized oral mucosa (alveolar mucosal keratinocytes, AM) and non-keratinized oral mucosa near the tooth surface (sulcular keratinocytes, SK). (c) Single-cell RNA sequencing (scRNAseq) was performed on adult healthy gingival tissues from mice; the UMAP represents the subclustering of murine keratinocytes. (d) Using Louvain clustering, *Krt14+; Krt19+* cells were found in small proportion to differentiated keratinized mucosal cells. (e) The majority of cells are basal (*Krt14* with and without *Ki67*) and suprabasal keratinocytes (*Krt4* and *Krt1*). (d) Though *Krt19* was expressed as mRNA, *Krt19* was not detected at the protein level. (f) Looking at similar cell signatures discovered in humans, many markers are not specific, though there appears to be some expression of immune markers in *Krt19* (Cluster 7), including *Cxcl1, Cxcl16, Cxcl17,* and *Cxcl3*. Another cluster (6, pink box) expressed some interleukin inflammatory markers unique to cluster 7 (yellow box) that are shared in humans. Abbreviations: Molar (M); see Figure 1 legend. Scale bars: (b) 50 μm, (c) 100 μm. Illustration from (a) created with BioRender. com.

**Extended Data 6.**
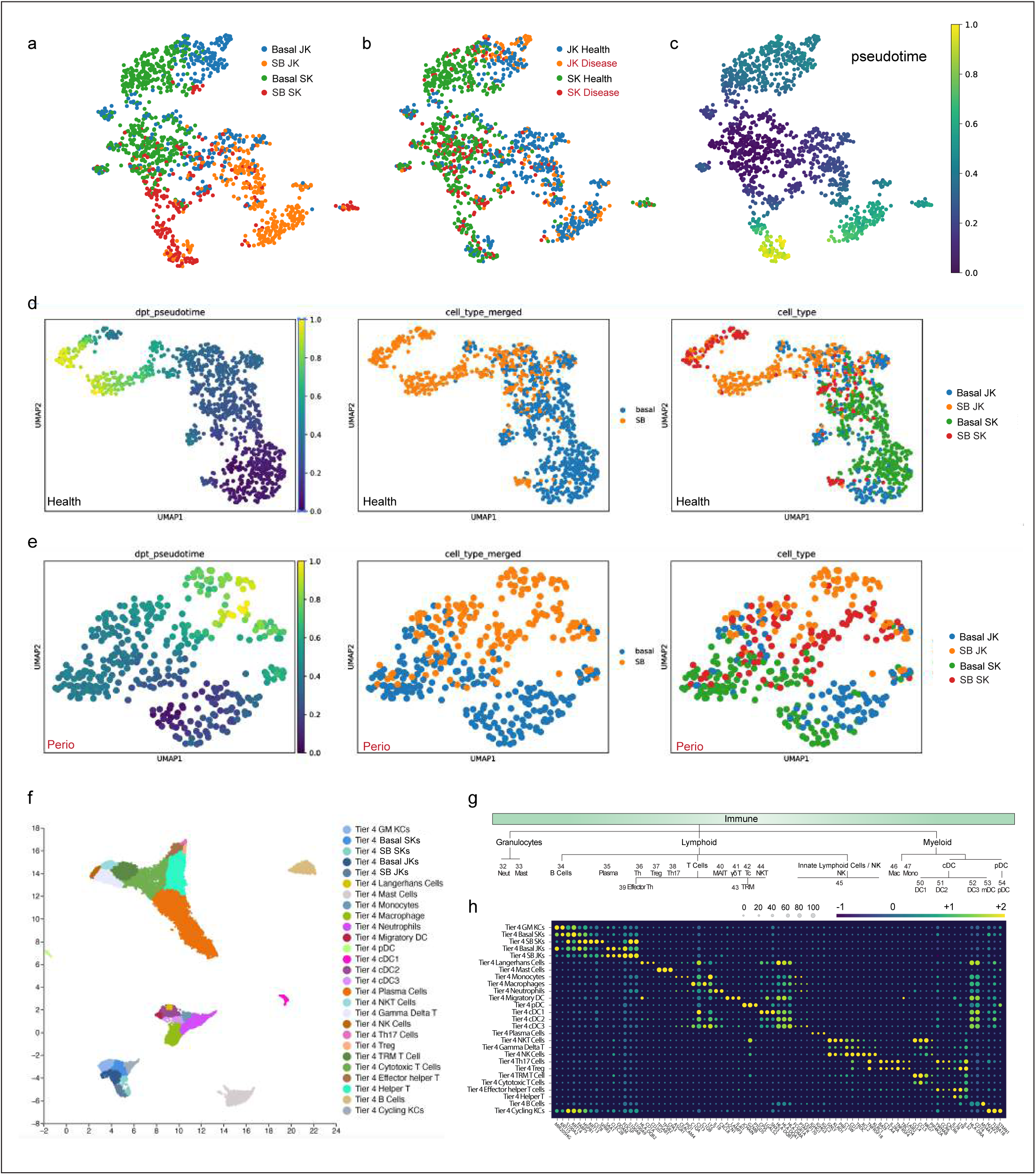
Tier 4 annotation of epithelial and immune cell types of the periodontium. (a-c) *KRT19*-high JK, SK, and GM keratinocytes (KCs) were subclustered for further annotation using PAGA. (d-e) PAGA broken out for health (d) and periodontitis (e). (f-g) Using CellTypist, we created another Tier 4 UMAP (f) and refined the first draft annotation of innate and adaptive immune cell subpopulations (g) that were included in the CellChat analysis74 from Figure 3. (h) Markers are included for each cell subpopulation. Abbreviations: Tissue Resident Memory T Cell (TRM T); also see Figure 1 legend.

**Extended Data 7.**
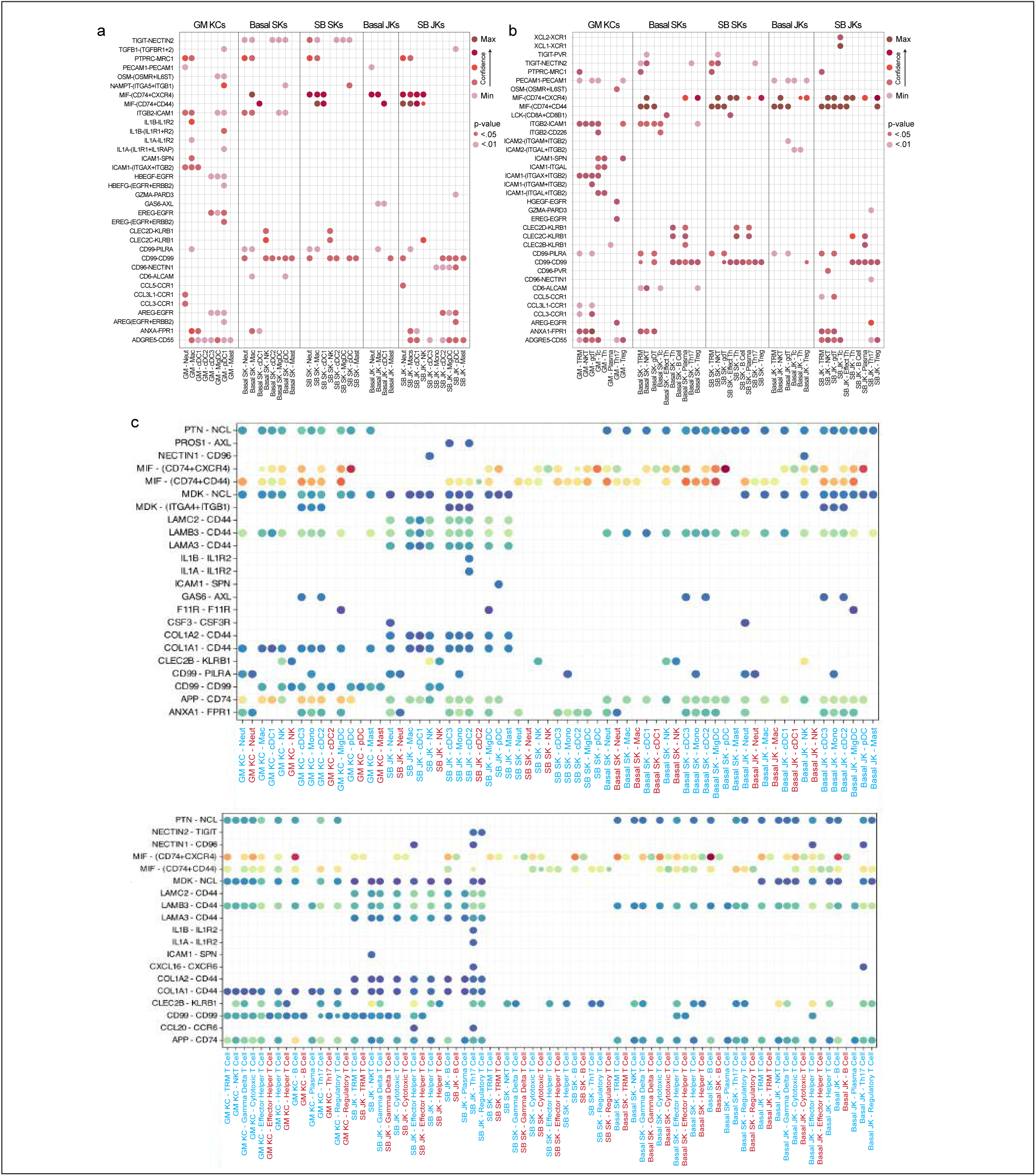
CellChat reveals up-and downregulated communication between cell subpopulations in periodontitis. (a-b) Significant pathway utilization at a single receptor-ligand level is plotted by confidence and p-value significance. These plots focus on GM, SK, and JK populations and reveal distinct differences in innate (a) and adaptive (b) signaling pathways between basal and differentiated KC populations. (c,d) Dot plots show downregulated receptor-ligand interactions by cell type and by disease state in innate (c) and adaptive (d) populations. Abbreviations: see Figure 1 legend.

**Extended Data 8.**
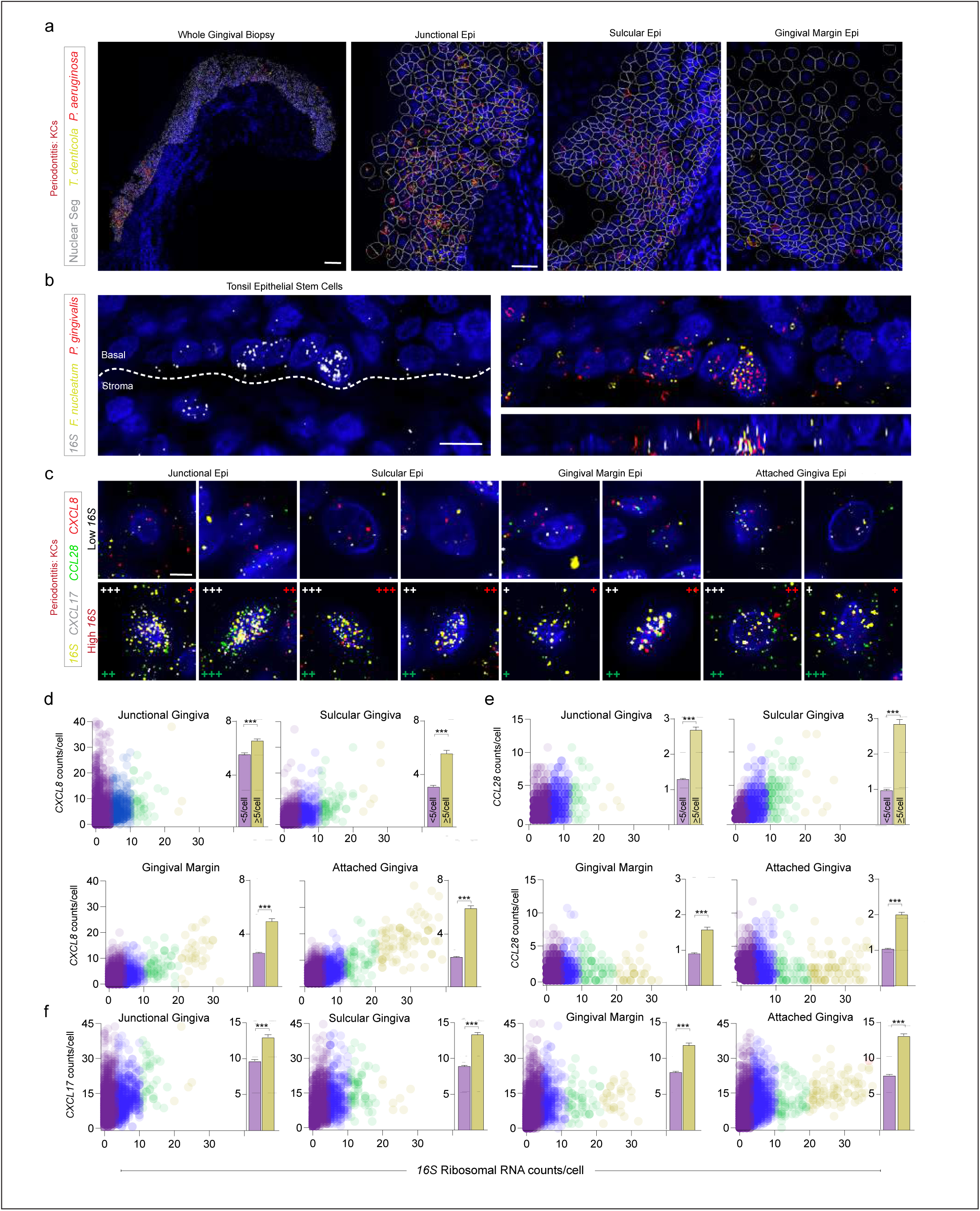
Polybacterial coinfection is not specific to gingiva but can be enriched in JKs. (a) Some periopathogens are enriched in JKs where the obligate anaerobes such as *T. denticola* survive in an oxygen-poor environment. Facultative anaerobes such as *P. aeruginosa* can tolerate JK, SK, ad GM environments, making coinfection dependent on niche environmental conditions. (b) Evidence for polybacterial infection in tonsil epithelial stem cells from classically defined “periopathogens” *P. gingivalis and F. nucleatum*. (c-f) Further evidence at a single-cell level that *CCL28, CXCL8,* and *CXCL17* are positively correlated with *16S* levels (c), independent of keratinocyte spatial localization. Quantification phenotypes in Figure 8 reveal statistically significant associations between bacterial burden and *CXCL8* (d), *CCL28* (e), and *CXCL17* (f). Abbreviations: see Figure 1 legend. Scale bars: (a,c) 10 μm; (b) 100 μm (insets: 25 μm); p<0.05, paired Student’s T-test (d-f).

## METHODS

### HUMAN INTEGRATED PERIODONTAL META-ATLAS GENERATION AND ANALYSIS

Human single-cell data reprocessing: Raw fastq files for the previously published single-cell RNA sequencing projects were downloaded and processed using scripts available here: https://github.com/cellgeni/ reprocess_public_10x. Briefly, series metadata was collected using the GEO soft family file. Following this, ENA web API was used to obtain information about the format in which raw data is available for every run (SRR/ERR), as well as to infer the sample-to-run relationships. Raw read files were then downloaded in one of the three formats: 1) SRA read archive; 2) submitter-provided 10X BAM file; 3) gzipped paired-end fastq files. SRA archives were converted to fastq using fastq-dump utility from NCBI SRA tools v2.11.0 using “-F --split-files” options. BAM files were converted to fastq using 10X bamtofastq utility v1.3.2. Following this, raw reads were mapped and quantified using the STARsolo algorithm. STAR version 2.7.10a_alpha_220818 compiled from source files with the “-msse4.2” flag was used for all samples. Wrapper scripts documented in https://github.com/cellgeni/STARsolo/ were used to auto-detect 10x kit versions, appropriate whitelists, and other relevant sample characteristics. The human reference genome and annotation exactly matching Cell Ranger 2020-A was prepared as described by 10x Genomics: https://support.10xgenomics.com/single-cell-gene-expression/software/release-notes/build#header. For 10x samples, the STARsolo command was optimized to generate the results maximally like Cell Ranger v6. Namely, “--soloUMIdedup 1MM_CR --soloCBmatchWLtype 1MM_multi_Nbase_ pseudocounts --soloUMIfiltering MultiGeneUMI_CR --clipAdapterType CellRanger4 --outFilterScoreMin 30” were used to specify UMI collapsing, barcode collapsing, and read clipping algorithms. For paired-end 5’ 10x samples, options “--soloBarcodeMate 1 --clip5pNbases 39 0” were used to clip the adapter and perform paired-end alignment. For cell filtering, the EmptyDrops algorithm employed in Cell Ranger v4 and above was invoked using “--soloCellFilter EmptyDrops_CR” options. Options “--soloFeatures Gene GeneFull Velocyto” were used to generate both exon-only and full-length (pre-mRNA) gene counts, as well as RNA velocity output matrices.

Cellenics® database generation and subclustering: The single-cell RNA-seq dataset was processed, analyzed and visualized using the Cellenics® community instance (https://scp.biomage.net/) hosted by Biomage (https://biomage.net/). Pre-filtered count matrices were uploaded to Cellenics®. Barcodes were then filtered in a series of four sequentially applied steps. Barcodes with less than 500 UMIs were filtered out. Dead and dying cells were removed by filtering out barcodes with a percentage of mitochondrial reads above 15%. To filter outliers, a robust linear model was fitted to the relationship between the number of genes with at least one count and the number of UMIs of each barcode using the MASS package (v. 7.3-56)^75^. The expected number of genes was predicted for each barcode using the fitted model with a tolerance level of 1 - alpha, where alpha is 1 divided by the number of droplets in each sample. Droplets outside the upper and lower boundaries of the prediction interval were filtered out. Finally, the probability of droplets containing more than one cell was calculated using the scDblFinder R package v. 1.11.3^76^. Barcodes with a doublet score greater than 0.5 were filtered out. After filtering, each sample contained between 300 and 8000 high-quality barcodes and was input into the integration pipeline. In the first integration step, data was log-normalized, and the top 2000 highly variable genes were selected based on the variance stabilizing transformation (VST) method. Principal-component analysis (PCA) was performed, and the top 40 principal components, explaining 95.65% of the total variance, were used for batch correction with the Harmony R package^71^. Clustering was performed using Seurat’s implementation of the Louvain method. To visualize results, a Uniform Manifold Approximation and Projection (UMAP) embedding was calculated, using Seurat’s wrapper around the UMAP package^77^. To identify cluster-specific marker genes cells of each cluster were compared to all other cells using the presto package implementation of the Wilcoxon rank-sum test^71^. Keratinocytes were subset from the full experiment by extracting manually annotated barcodes and filtering the Seurat object. The subset samples were subsequently input into the Biomage-hosted instance of Cellenics®. Filtering steps were disabled since the data was already filtered. The data was subjected to the same integration pipeline as the full experiment. All cells were manually annotated using available literature and CellTypist^74^.

Transfer of Cellenics® data to cellxgene: Annotated cell-level data was downloaded from Cellenics in the form of an rds file containing a Seurat object. The data was converted by exporting count matrices and metadata from R and loading them using Scanpy version 1.9.3 (https://scanpy.readthedocs.io/). Additional metadata (e.g., age, sex, self-reported ethnicity) from the original datasets were matched to the closest entries in the respective ontology, per cellxgene contribution guidelines (https://cellxgene.cziscience.com/docs/032__Contribute%20and%20Publish%20Data).

DEG Analysis using g:Profiler^78^: Differentially expressed gene (DEG) lists were generated in Cellenics® and exported as .csv files and uploaded to the g:Profiler website (https://biit.cs.ut.ee/gprofiler/gost). g:Profiler is part of the ELIXIR Recommended Interoperability Resources that support FAIR principles. A complete list of those resources can be found: https://elixir-europe.org/platforms/interoperability/rirs. g:Profiler assesses Gene Ontology and pathways from KEGG Reactome and WikiPathways DEGs were uploaded to the query section and were first analyzed using g:GOSt multi-query Manhattan plots. These data were further analyzed for the results tab (GO:MF, GO:CC, GO:BP, KEGG, REAC, TF, MIRNA, HPA, CORUM, HP, WP). Data from the Extended Data figures are an incomplete display of all the g:Profiler data. DEGs are included in Supplementary Table 1 for further analysis.

CellPhoneDB^79^ and CellChat80: The total number of ligand-receptor interactions between tier 3 cell types was calculated for healthy and periodontal disease using CellPhoneDB (version 3.1.0). The tier 3 annotated AnnData object was subsetted and separate AnnData objects saved for healthy and periodontal disease, respectively. Metadata tables containing the cell barcodes as indices were also exported. CellPhoneDB was then run as follows: cellphonedb method statistical_analysis metadata. tsv AnnData.h5ad --iterations=10 --counts-data hgnc_symbol threads=2. The CellPhoneDB results were filtered by removing those interactions with a P value > 0.05. Results were visualized using a modified form of CellPhoneDB’s plot_cpdb_heatmap function to allow for re-ordering of cell types. Cell-cell interactions between receptors in tier 4 keratinocyte subtypes and ligands in innate and adaptive immune subtypes were further explored using the R package CellChat (version 1.6.1 using the cell-cell interaction database). The tier 4 annotated AnnData object was subsetted and separate expression matrices exported for healthy and periodontal disease, together with their respective metadata tables. These were used to create Seurat objects for healthy and periodontal disease, which served as input to CellChat. Analyses were performed using the log-transformed normalized gene counts with default parameters and using the human CellChatDB. Cell type composition differences were accounted for when calculating communication probabilities. Data from healthy and disease were compared to identify significant changes.

Partition-based graph abstraction (PAGA) plots81: The keratinocyte subset count matrices were imported into Scanpy version 1.9.3^81^ to conduct quality control, normalization, and log-transformation of the data to control for variability in sequencing depth across cells. To minimize the potential batch effects across the four datasets, a batch correction technique was applied using the Python package HarmonyPy version 0.0.9^71^, with the ‘sample ID’ serving as the batch key. PAGA graphs were constructed using Scanpy’s implementation. These graphs were used to explore the relationships between different clusters of cells and to understand the potential developmental trajectories. The coordinates for Uniform Manifold Approximation and Projection (UMAP)^82^ were then calculated with the PAGA graph as the initial position, allowing for a visualization that is coherent with the topology of the PAGA graph. To better understand the developmental progression of cells along these trajectories, pseudotimes were estimated by diffusion pseudotime (DPT) analysis^83^ over the PAGA graphs. The DPT is a measure of the transcriptional progression of cells along a trajectory, starting from root cells that were manually selected. Heatmaps were created to visualize gene expression changes along the trajectories, with manually selected start and endpoints, using both Scanpy and seaborn^84^. To smooth the plots and reduce noise, a moving average of the expression values was used, with a window size of 50 data points along pseudotime. The clustering of the genes in the heatmaps was performed using Ward’s method.

Mouse single-cell RNA library preparation, sequencing, processing, and analysis of data: All the necessary animal procedures were followed according to the UK law, Animals Scientific Procedures Act 1986. The experiments were covered by the necessary project licenses under the Home Office and Queen Mary University of London’s institution’s Animal Welfare & Ethical Review Body (AWERB). The mouse tissues were obtained at Queen Mary University of London, Barts & The London School of Medicine and Dentistry. Mice from both genders were maintained on the C57BL/6 N genetic background and were housed under a 12-hour light/12- hour dark cycle, at temperatures of 20–24°C with 45–65% humidity. Single-cell suspensions of gingival tissue were obtained from P28 mice, sacrificed by cervical dislocation. Three biological replicates were pooled together to give one single sample for sequencing. Both males and females were used. Fresh gingival tissues were processed immediately after dissection, cut into smaller pieces in a sterile petri dish with RPMI medium (#11875093, Sigma) and digested for 30 min at 37°C under agitation using the Miltenyi Mouse-Tumor Dissociation kit (#130-096-730). The resulting cell suspension was consecutively filtered through 100 µm and 70 µm cell strainers and cells were collected by centrifugation. The viability of the cell suspension was determined using a Luna-FL automated cell counter (Logos Biosystems). Single-cell cDNA library was prepared using the 10X Genomics Chromium Single-cell 3’ kit (v3.1 Chemistry Dual Index). The prepared libraries were sequenced on Illumina® NovaSeq™6000 (2×150 bp) with a targeted sequencing depth of ∼30,000 reads/cell. The cell ranger-6.0.1 pipeline was used for processing the scRNAseq data files before analysis according to the instructions provided by 10x Genomics. Briefly, base call files obtained from each of the HiSeq2500 flow cells used were demultiplexed by calling the ‘cellranger mkfastq’. The resulting FASTQ files were aligned to the mouse reference genome (GRCm38/ mm10), filtered, and had barcodes and unique molecular identifiers counted and count files generated for each sample. The raw count matrix output from CellRanger was then processed by the ambient RNA removal tool CellBender^85^, giving an output filtered count matrix file. This was used for subsequent preprocessing and data analysis using Python package 3.8.13 with the Scanpy pipeline. For basic filtering of our data, we filtered out cells expressing less than 200 genes and less than 100 counts. We filtered out genes expressed in less than three cells and with less than 10 counts. Cells were filtered out by applying the following thresholds: 1) more than 20% mitochondrial reads; 2) ribosomal reads lower or higher than the 5th and 95th percentile; 3) more than 1% of hemoglobin reads and 4) total reads lower than 700 and higher than 50000. Scrublet, a doublet removal tool was applied to further remove predicted doublets. To ensure that the data is comparable among cells, we normalized the number of counts per cell to 10,000 reads per cell. Data were then log-transformed for downstream analysis and visualization. The cell cycle stage was predicted using the sc.tl.score_genes_cell_cycle tool4. We regressed out cell-to-cell variations driven by mitochondrial, ribosomal, and cell-cycle gene expression and the total number of detected molecules. We then scaled the data to unit variance. The neighborhood graph of cells was computed using PCA presentation (n PCs = 40, n neighbors = 10). The graph was embedded in two dimensions using Uniform Manifold Approximation and Projection (UMAP) as suggested by Scanpy developers. Clusters of cell types were defined by the Leiden method for community detection on the generated UMAP graph at a resolution of 0.1. Epithelial clusters were used for the second-level clustering. The respective cell types were identified upon annotation of clusters from first-level clustering. The cluster-specific barcodes were retrieved as a list, which was used to select the cells of interest from the filtered count matrix on a separate Jupyter notebook and were re-analyzed separately. Epithelial cells were filtered and analyzed as previously described, and clustered at resolution 0.5 using Louvain.

Single-cell Analysis of Host-Microbiome Interactions (SAHMI)^86^: The standard Kraken2 database (version 2.1.3) was downloaded. To avoid overlooking potential oral microbes, genomes from the Human Oral Microbiome Database (HOMD) (PMID 20624719) not present in the standard database (n=1,502 taxIDs) were also downloaded, and this custom database was built using kraken2-build^87^ with default parameters. Reads from were taxonomically classified using Kraken 2, with “–use-names” and “–report-minimizer-data” (Kraken2Uniq) but otherwise default parameters. True positives from Kraken2 results were identified using barcode level denoising from the SAHMI pipeline and rRNA enrichment. First, barcode denoising was performed. True taxa were identified by performing Spearman correlations between the number of unique and total k-mers across barcodes in each sample. Taxa found to significantly correlate (p-value < 0.05) in at least one sample were retained. For all retained taxa, genomic contigs belonging to the taxa were extracted from the Kraken2 database and the reads that were classified to that specific taxa were then mapped to those genomic contigs using bowtie2 (version 2.2.5)^88^ with default parameters. Additionally, rRNAs were annotated along the extracted genomic contigs using barrnap (version 0.8) with default parameters. BEDTools (version 2.30.0)^89^ coverage was used to count the number of aligned reads overlapping annotated rRNAs. We then calculated the fold enrichment of reads across rRNAs relative to the entire genome, normalized by rRNA and genome length, respectively. Taxa found to contain at least a 5-fold enrichment in rRNA sequences relative to the whole genome, which is expected for bacterial transcriptomics that is not rRNA depleted, were retained. From the human host reads, we previously identified which barcodes corresponded to which cell types. Because the reads that are classified to microbial taxa also contain these same barcodes, we then assigned cell types to the microbial reads. To calculate the relative abundance of taxa in each cell type, we divided the total number of reads classified to those taxa with a barcode assigned to that cell type by the total number of reads in the sample assigned to that cell type.

### SPATIAL VALIDATION AND ANALYSIS OF KERATINOCYTES

Mouse husbandry: All mice (Extended Data 5e) were bred and maintained in an AAALAC-certified animal facility under an IACUC-approved protocol (ID#20-041.0-B) at the University of North Carolina at Chapel Hill. Each animal was determined to have a healthy body score of at least 3 and had not previously been included in any other panel. Mice were euthanized in accordance with the Panel on Euthanasia of the American Veterinary Medical Association.

Tissue preparation, mounting, and sectioning: Deidentified human gingival tissues were acquired from discarded routine third molar extractions or from gingival biopsies (UPenn to LOCI; IRB #844933; MTA #68494). Immediately after extraction, tissues attached to third molars were placed in a 10% solution of NBF and fixed for a minimum of 24 h in a 4˚C refrigerator. After fixation, the tissues were washed twice in 1X PBS before being placed in 70% EtOH in a 4 ˚C refrigerator until they were ready to be mounted. Tissues were embedded in paraffin blocks using a Leica system and stored in a 4 ˚C refrigerator until sectioning using RNAse precautions on a Leica system. Formalin-fixed, paraffin-embedded (FFPE) human gingival tissue on SuperFrost Plus slides was heated to 60 ˚C on a slide warmer for 30 min. Following deparaffinization for 10 min using HistoChoice Clearing Agent, the tissues were rehydrated using a series of ethanol solutions (100%, 90%, 70%, 50%, and 30% EtOH in nuclease-free water) for 10 min (for 100%) and 5 min each, followed by 2 x 5 min in 100% nuclease-free water. During rehydration, 50 mL of a 1X solution of AR9 buffer in nuclease-free water was prepared and added to a Coplin jar. Following rehydration, the slides were added to the 1X AR9 buffer and covered with aluminum foil. Samples were antigen retrieved in a pressure cooker for 15 min at low pressure. Following antigen retrieval, the Coplin jar was removed from the pressure cooker and cooled for at least 30 min. The slide was then soaked in nuclease-free water for 30s, followed by soaking in 100% EtOH for 3 min, both in Coplin jars.

Immunohistochemistry (IHC) on Human or Murine Tissues: Blocking solution was prepared using reagents in Table 1. The sample underwent antigen retrieval as described. A pap-pen was used to draw a hydrophobic barrier around the sample and allowed to dry for 5 min. The slide was then placed in a humidity chamber, and the sample was washed with 1X PBS for 2 x 5 min using a Pipetman (Gibco), then blocked using the blocking solution for 1 h. During blocking, an antibody cocktail using primary anti-human antibodies (Table below) in blocking solution dilution was prepared. Following the removal of the blocking solution, this antibody cocktail was added to samples, which were stored overnight in a 4˚C refrigerator. The next day, a secondary antibody cocktail was prepared with AlexaFluor 488 (AF488), Rhodamine Red-X (RRX), and Cyanine 5 (Cy5; Table below for concentrations). The primary antibody cocktail solution was removed, and the samples were washed with 1X PBS for 2 x 5 min. The secondary antibody cocktail was added to samples and left to hybridize for 2h at room temperature. This cocktail was then removed, and the samples were washed with 1X PBS for 2 x 5 min. Then, a solution of DAPI (1:1000 in 1X PBS) was added to the sample for 5 min. This solution was removed, and the sample was washed with 1X PBS for 2 x 5 min before mounting with ProLong Gold Antifade. Imaging was performed using a Leica DMi8 with THUNDER Imager (Leica Microsystems) using a 20X or 40X objective.

| Primary antibody | Host | Species | Clone | Dilution | Vendor |
| --- | --- | --- | --- | --- | --- |
| Keratin 14 (KRT14) | Ck | Hu | Poly9060 | 1:500 | BioLegend |
| Keratin 19 (KRT19) | Rb | Hu, Ms | EP1590Y | 1:500 | Abcam |
| Ki67 | Rt | Hu, Ms, Rt | SolA15 | 1:500 | Invitrogen |

| Target | Host | Fluorescence | Dilution | Vendor |
| --- | --- | --- | --- | --- |
| Rb | Dk | AlexaFluor 488 | 1:1000 | Jackson |
| Rt | Dk | Rhodamine Red-X | 1:500 | Jackson |
| Ck | Dk | Cyanine 5 | 1:400 | Jackson |

RNAscope HiPlex 12 V1 or V2 on Human Tissues: All reagents in this section were purchased and used as received from ACD unless otherwise noted. The sample underwent deparaffinization and rehydration as described. 1 drop of RNAscope hydrogen peroxide was added to the slides, and the samples were left for 10 min at room temperature. The H2O2 was tapped off the slides, and the samples were antigen retrieved and dried as described. During this time, the HybEz oven was turned on and set to RNAscope (40˚C). The hydration paper was wetted with nuclease-free water to prepare the humidity chamber in the slide tray. An Immedge pen was used to draw a tight hydrophobic barrier around the tissues, then dried at room temperature for 5 min. The slides were then placed in the slide holder. One drop of Protease IV reagent was added to each contained region. The slide holder was placed in the tray, and the tray was placed in the HybEz oven for 30 min. During this time, 1X RNAscope wash buffer was prepared in nuclease-free water. The RNAscope hybridization solutions were prepared by adding 1 µL of T probe (Table below) to 100 µL of probe diluent. The tray was removed, and the slide carrier was immersed in the wash buffer for 2 x 2 min. The carrier was removed and dried, and a paper towel was used to dry the area around the barrier. Then, 20 µL of hybridization solution or 1 drop of positive or negative control probe was added to each spot. The slide holder was replaced in the tray, and the tray was placed in the HybEz oven for 2 h. After 2h, the slide holder was washed in wash buffer for 2 x 2 min. Signal amplifiers were added to the samples by hybridization of AMP1, AMP2, and AMP3 for 30 min, 30 min, and 15 min, respectively with washing in-between steps. After signal amplifiers, T1-T4 fluorophores were added to each spot, with 15 min hybridization and washing. Then, RNAscope DAPI was added to each sample for 30s. Following this, the DAPI was tapped off the slides, which were immediately mounted with Prolong Gold Antifade. Imaging was performed using a Leica DMi8 with THUNDER Imager (Leica Microsystems) using a 40X or 63X objective. After imaging acquisition had been completed for T1-T4 probes, the sample necessitated the removal of the first probes for imaging of the second probes. After completed imaging, slides containing samples were placed in 4X saturated sodium citrate (SSC) for 30 min. During this time, an ampule of RNAscope cleaving solution was opened, and the contents were added to 10 mL of 4X SSC. The slides were left immersed in 4X SSC until the cover glass could be gently removed, after which the slides were added to the slide holder and one drop of cleaving solution was added to each region containing the sample. The slide holder was loaded into the tray, and the tray was loaded into the HybEz oven for 15 min. Then, the slide holder was washed in 0.5% Tween for 2 x 2 min. This process was repeated. Following this, T5-T8 probes were added to the sample in the same manner, and the sample was imaged. Once the imaging of T5-T8 probes was completed, their reporters were cleaved, and the T9-T12 probes were hybridized and imaged. For HiPlex V2, Protease III was used instead of Protease IV. Additionally, for HiPlex V2, between the AMP3 and the addition of T1-T4, the addition of FFPE reagent was required as follows. To 100 µL of 4X SSC was added 2.5 µL of FFPE reagent, resulting in a 1:40 solution of FFPE reagent. Following washing slides using 1X wash buffer, the FFPE reagent was added to each slide and incubated in the HybEz Oven for 30 min. Following this, the slide holder was removed from the tray and immersed in 1X wash buffer before proceeding to fluorophore addition.

| Target | Channel | Fluorescence | NBCI ref sequence | Chemistry |
| --- | --- | --- | --- | --- |
| KRT19 | T1 | 488 | NM_002276 | V1 |
| CCL28 | T1 | 488 | NM_148672.2 | V2 |
| Polr2a (+ control) | T1 | 488 | NM_000937.4 | V1 |
| LAMB4 | T2 | 550 | NM_001318048.1 | V1 |
| CXCL17 | T2 | 550 | NM_198477.2 | V2 |
| P. gingivalis | T2 | 550 | NR_040838.1 | V2 |
| PPIB (+ control) | T2 | 550 | NM_000942.4 | V1 |
| CXCL8 | T3 | 647 | NM_000584.3 | V2 |
| F. nucleatum | T3 | 647 | NR_026083.1 | V2 |
| UBC (+ control) | T3 | 647 | NM_021009 | V1 |
| CXCL14 | T4 | 750 | NM_004887.4 | V1 |
| EB-16S | T4 | 750 | * | V2 |
| HPRT1 (+ control) | T4 | 750 | NM_000194.2 | V1 |
| SAA1 | T5 | 488 | NM_000331.6 | V1 |
| IL36G | T5 | 488 | NM_001278568.1 | V2 |
| COL7A1 | T6 | 550 | NM_000094.3 | V1 |
| RHOV | T6 | 550 | NM_133639.3 | V1 |
| IL1B | T6 | 550 | NM_000576.3 | V2 |
| T. denticola | T6 | 550 | NC_002967.9:610211-611726 | V2 |
| NPPC | T7 | 647 | NM_024409.2 | V1 |
| CCL20 | T7 | 647 | NM_001130046.1 | V2 |
| P. aeruginosa | T7 | 647 | ENA\|AB680164.1 | V2 |
| ODAM | T8 | 750 | NM_001385579.1 | V1 |
| PAPPA | T8 | 750 | NM_002581.5 | V1 |
| IL1A | T8 | 750 | NM_000575.3 | V2 |
| SFRP1 | T9 | 488 | NM_003012.4 | V1 |
| TNFSF15 | T9 | 488 | NM_005118.3 | V2 |
| IL18 | T10 | 550 | NM_001562.3 | V1 |
| IL6 | T10 | 550 | NM_000600.3 | V2 |
| RHCG | T11 | 647 | NM_001321041.2 | V1 |
| CXCL3 | T11 | 647 | NM_002090.3 | V2 |
| NEAT1 | T12 | 750 | NR_131012.1 | V1 |
| CXCL1 | T12 | 750 | NM_001511.4x | V2 |
| FDCSP | T11 | 647 | NM_152997.3 | V1 |
| LGR6 | T6 | 550 | NM_001017404.1 | V1 |
| dapB (- control) | T1-T4 | various | EF191515 | V1 |
| SOX6 | T4/T12 | 750 | NM_017508.2 | V1 |

PhenoCycler-Fusion (PCF) on Human Tissues: All reagents in this section were purchased and used as received from Akoya Biosciences unless otherwise noted. Samples underwent deparaffinization, rehydration, and antigen retrieval as described above. Following sample immersion in EtOH, the sample was immersed in Akoya Hydration Buffer for 2 min, followed by Akoya Staining Buffer for 20 min. While the sample cooled to rt, the antibody cocktail was prepared. To 362 µL of Akoya Staining Buffer was added 9.5 µL of N, J, G, and S blockers. Then, 150 µL of blocking solution was pipetted into a 1.5 mL vial, and 1 µL of barcoded antibody was added to the vial such that the final volume of antibody blocking solution was 190 µL. After immersion in Staining Buffer, the slide was removed, the back and area around the sample were wiped dry, and the slide was added to a humidity chamber. As a modification to the manufacturer’s instructions, the antibody blocking solution was added to the sample, and the humidity chamber was placed in a 4˚C refrigerator overnight. After removal of the blocking solution, the slide was placed in staining buffer for 2 min, followed by post-stain fixing solution (10% PFA in staining buffer) for 10 min. Following 3 x 2 min washing in 1X PBS, the slide was immersed in ice-cold MeOH for 5 min. While the slide was immersed, the final fixative solution was prepared by adding 1 vial of fixative to 1 mL of staining buffer. The slide was removed from MeOH and placed in the humidity chamber, and 200 µL of the final fixative solution was added to the sample. This was left in place for 20 min. Then, the final fixative solution was removed, and the slide was washed in 3 x 2 min in 1X PBS. To convert the slide into a flow cell for use in the PCF experiment, the back of an Akoya flow cell top was removed, and the top was placed adhesive face up in the Akoya-provided impressing device. The slide was removed from the 1X PBS, and the edges around the slide that matched where the top of flow cell adhesive would adhere were dried using a micro-squeegee toolkit (Essential Bangdi). Then, the slide which formed the bottom of the flow cell was placed sample-side down on the top of the flow cell without applying pressure to the adhesive. The tray of the impressing device was inserted into the device, and the lever was gently pulled to adhere to the top and bottom of the flow cell. After 30 s, the lever was depressed, the tray was pulled out, and the flow cell was removed. This flow cell was placed in 1X PCF buffer without buffer additive for a minimum of 10 min before any PCF experiment to allow for improved adhesion between the top and bottom of the flow cell. To prepare the PCF reporter wells, a 15 mL Falcon tube was first wrapped with aluminum foil. To this Falcon tube was added 6.1 mL of nuclease-free water, 675 µL 10X PCF buffer, 450 µL PCF assay reagent, and 4.5 µL of in-house prepared concentrated DAPI such that the final DAPI concentration was 1:1000. Then, this reporter stock solution was pipetted to 18 amber vials, with the volume in each vial being 235 µL. To each vial was added 5 µL of reporter per cycle. The total volume per vial was either 245 µL for a cycle with 2 reporters or 250 µL for a cycle with 3 reporters; to optimize reagents and reporters, no cycles contained only 1 reporter. Only one criterion was used to create a cycle: each cycle could contain a maximum 3 reporters, corresponding to 1 of Atto550, AlexaFluor 647, and AlexaFluor 750 (where appropriate; see below for more information). A separate pipet tip was used to pipet the contents of each amber vial to a 96-well plate. DAPI-containing vials were pipetted into a well in the H-row, whereas vials containing reporters were pipetted into wells in other rows. Once all wells were filled, Akoya-provided foil was used to seal the wells. Imaging was performed using a PhenoImager Fusion connected to a PhenoCycler i.e., the PhenoCycler Fusion system (Akoya BioSciences) using a 20X objective (Olympus). Requisite solutions for this instrument include ACS-grade DMSO (Fisher Chemical), nuclease-free water, and 1X PCF buffer with buffer additive, the latter of which was prepared by adding 100 mL of 10X PCF buffer and 100 mL of buffer additive to 800 mL of nuclease-free water.

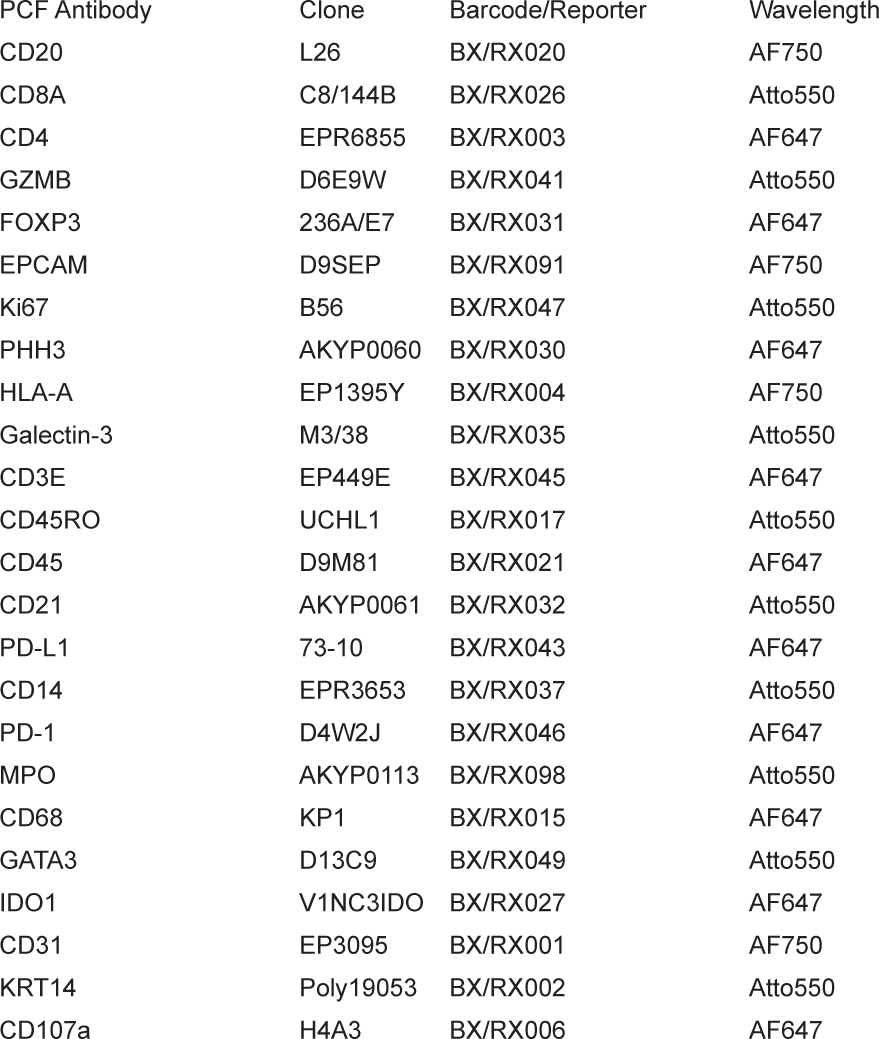
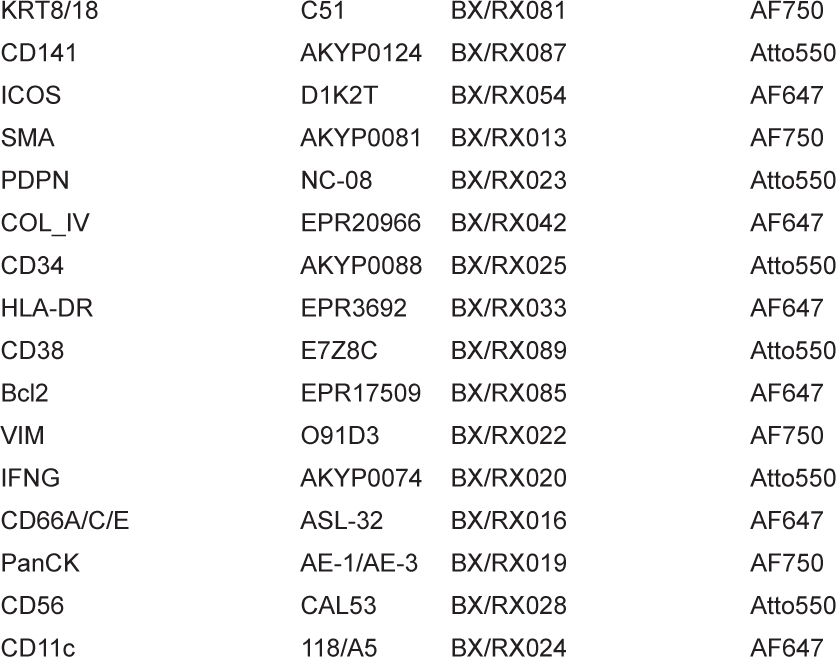

### HUMAN PRIMARY CELL CULTURE AND ANALYSIS

Human gingival keratinocyte (HGK) culture passaging, cryopreservation, and fixation: All reagents in this section were purchased and used as received from ATCC (Extended Data 4) or Lifeline Cell Technologies (Figure 5). DermaLife K Keratinocyte Medium (HGK medium), the keratinocyte growth kit, and a 6-well plate (ThermoFisher Scientific) were brought into a biosafety cabinet using aseptic techniques. The keratinocyte growth kit reagents were added to the dermal basal cell medium). To 3 of the wells was added 1.5 mL of warmed media. The HGKs (passage (P)2: Lifeline FC-0094 lot #05390; P2 ATCC PCS-200-014 lot #80523333) were thawed and aliquoted into 100 µL portions in cryovials (ThermoFisher). One of the 100 µL vials was diluted to 1.5 mL by adding media; then, 500 µL of this diluted media was added to each well, and the 6-well plate was placed in a tissue cabinet at 5% CO2 and 37 °C. After 24 h, the media was removed and replaced. Cells were grown in the well until reaching 70% confluence, at which point they were passaged using 0.05% trypsin EDTA, neutralized, and pelleted using a centrifuge, resulting in P3 HGKs. Some cells were re-plated in HGK medium and followed the same procedure until P4 and P5 HGKs were obtained. Cells that were not plated or re-plated were cryopreserved. P2 HGKs were cryopreserved using the solution in which they were delivered. For in-house passaged HGKs, the excess trypsin/neutralizing solution was removed, and the HGKs were then resuspended in 1 mL of Frostalife (Lifeline). The cell suspensions were aliquoted into 1 mL cryovials, which were then placed into a Nalgene Freezing Container that was pre-loaded with 250 mL of isopropanol. The cells were cooled to -80°C in an ultra-low-temperature freezer for at least 2 h before immersion in liquid nitrogen. Instead of cryopreservation, some cells were fixed for downstream analysis. In a biosafety cabinet, bacterial LPS from P. gingivalis (Invivogen) was added to endotoxin-free water (Invivogen). Serial dilution was used to prepare a solution of 20 µg/mL of P. gingivalis LPS in HGK medium (LPS medium). P3 HGKs were plated in a 6-well plate and grown until a minimum of 40% confluency. Then, the HGK medium was removed, and 2 mL of LPS medium was added to three of the plates; the other three plates were filled with standard HGK medium. The cells were grown for 48h before the medium was removed. The cells were passaged and either cryopreserved or fixed. Following centrifugation and removal of the excess solution, the cells were suspended in 4% paraformaldehyde (PFA) for a minimum of 24h. Following fixation, the PFA solution was removed, and the cells were resuspended in 70% ethanol (EtOH) in water. These were stored in a 4°C refrigerator until future use. SuperFrost Plus (Fisher Scientific) slides were immersed in 0.1% poly-L-lysine solution in a Coplin jar for a minimum of 24 h. The slides were rinsed and dried at 37 °C for a minimum of 10 min before use. Cell suspensions were added to 1.5 mL Eppendorf tubes and diluted to 500 µL using 70% EtOH. Cytospin funnels were rinsed by adding an uncoated slide to the Cytospin clip and adding 70% EtOH to the funnel. The rotor was spun at 1600 RPM for 15 min, resulting in the evaporation of EtOH. Then, a poly-L-lysine slide was added to the Cytospin clip. The funnel was charged with the appropriate cell suspension in 70% EtOH and spun at 1600 RPM for 15 min to obtain cells in a circular spot on the slide. Multiple funnels were used, with rearrangement of the slide and funnels, to obtain 4 spots per microscope slide. Cells were processed for immunohistochemistry and RNA in situ hybridization (HiPlex 12) as outlined in the above sections.

Mycotesting was performed following manufacturer instructions using a MycoAlert Plus sample kit (Lonza) on a Tecan plate reader. Control testing was performed on Lonza Control sample and deionized water from the Millipore (MilliporeSigma). Samples tested included sterile media, media collected prior to passaging cells to P5, and media collected from LPS-challenged and unchallenged cells prior to passaging cells to P4.

### STATISTICAL METHODS

General Methods: All non-sequencing-based data were analyzed in Fiji, QuPath, and/or Prism 9. The selection of statistical tests is described in the text and figure legends. The graphs supporting each figure and the extended data were generated using Prism 9 and a community instance of Cellenics® (hosted by Biomage) unless otherwise specified. Venn Diagrams were generated using http://www.interactivenn.net/.

Spatial Proteomic Cell Assignment and Analysis. The determination of marker positivity in each cell was based on comparing their intensity to predefined thresholds. We carefully selected thresholds for each marker, specifically tailored to each slide, to ensure accurate cell type classification. A marker was positive if its intensity exceeds its threshold on the slide. Conversely, if the intensity fell below the threshold, the marker was considered negative. By applying this criterion, each cell was associated with positive/negative signals for each marker. The assignment of cell types for individual cells was made using predefined cell type signatures consisting of multiple markers. Cells considered fibroblast/stroma (PanCK-, CD45-, and CD31-), vascular cells (CD31+), and epithelia (PanCK+) were removed from further analysis. Only cells that were identified as immune cells (CD45+, PanCK-, and CD31-) were kept in the downstream analysis. In most cells, a unique cell type was confidently assigned based on the presence of the positive markers consistent with its signature. However, in some instances, a cell may exhibit positive markers from more than one cell type. To resolve this ambiguous cell-type assignment, we implemented a deconvolution approach (adapted from Celesta90). The intent was to assign the most likely cell type to each cell in the presence of cell-type mixtures. As such, for each mixed cell type group, we extract a feature intensity submatrix consisting of those cells and the features relevant to the cell type mixtures they represent. We then utilize a Louvain clustering method91 implemented in the Seurat toolkit version 392 to re-cluster those cells. The clusters were then subject to cell type assignment based on their enriched markers. Dot plots were used to visualize the marker intensity exhibited across those clusters. The identity of each cluster was then determined by inspecting highly expressed markers within. To infer cellular interactions from the spatial transcriptomics data annotated with cell types, we employed the Squidpy library in Python93. Each cell on the slide was represented as a node in the cellular interaction graph. Edges connecting the nodes were created using Delaunay triangulation and were assumed to represent two interacting cells. To remove excessively long edges, i.e., unlikely cell-to-cell interactions, a 99th percentile distance threshold was applied to the edges. From the cellular interaction graph, immune cells located in the junctional and sulcular stroma were extracted for further analysis. Interaction matrices were constructed to quantify the number of edges shared between each immune cell type within the junctional and sulcular stroma, respectively. Each entry in the interaction matrices represented the number of edges shared between the respective pair of immune cell types. To investigate the variation in immune cell interactions between the junctional and sulcular stroma, focusing on the levels of interaction between each immune cell type, we computed the difference in their respective interaction matrices. Specifically, we subtracted the interaction matrix of the junctional stroma from that of the sulcular stroma. The resulting matrix represented the difference in immune cell type interactions between the two regions, where more positive values represented greater interaction in the sulcular stroma and more negative values represented greater interaction in the junctional stroma. This subtraction matrix was then plotted as a hierarchically clustered heatmap using the Seaborn library in Python. This process was performed for each of the present slides to visualize the differences in immune cell interactions between the sulcular and junctional stroma across the various diseased patients. The resulting matrices were averaged to provide an aggregated view of the variations in the type of cells interacting with the immune cell.

Quantification and Plotting of In Situ Hybridization. A multi-step approach to analyze RNA multiplex images was utilized. after the image acquisition was performed using Fiji (ImageJ). The acquired LIF. files were subsequently converted into 8-bit images, as an ome.tiff file (scripts available at GitHub). The RNA multiplex images were acquired at three different time points, each time utilizing four probe sets in the same section. To ensure accurate representation, the three images from the same patient were overlaid using the Warpy and image combiner tools73, generating the 12-plex image represented in different channels. Subsequently, the images were subjected to segmentation using a pretrained model based on Cellpose 2.094 (Figure 6-12 Plex ISH in vivo) or on StarDist72 (other analyses). The segmentation process was iteratively refined using images from 80 fluorescent images from post-mortem biopsies. The model was trained multiple times until achieving accurate delineation of cell expansion in both the basal and suprabasal layers. The gingival biopsy was segmented separately for each of the areas of interest, sulcus, gingival margin, and junctional epithelial. each area divided by basal and suprabasal layers. Following the nuclei-based cell segmentation, subcellular analysis was performed using QuPath, where the number of RNA spots within each cell was detected. The raw data was then exported, and the number of RNA spots was detected and quantified per cell and channel. Subsequently, the raw data was processed as an input file through log scripts to rank the most highly expressed RNA transcripts and extract the field-of-view (FOV) from the X and Y coordinates of each cell. Manual thresholding was applied to define the positivity of the transcripts. To analyze the RNA transcript data further, we transformed the matrix into a Seurat object. The data was treated, and raw clustering was performed in the UMAP projection, as well as PCA evaluation, for each of the twelve probes analyzed, adding the spatial information regarding RNA transcript expression in each patient sample, providing RNA expression patterns within the basal and suprabasal layers from each of the ROIs in the gingival tissue. Individual cell quantification (Extended Data 8) was performed using a similar process (segmentation, manually thresholded dot quantification) but was conducted independently to validate findings.

### CODE AVAILABILITY

Analysis notebooks are available at: http://github.com/LOCI/periodontitis, https://github.com/cellgeni/reprocess_public_10x, https://github.com/ cellgeni/STARsolo/, https://support.10xgenomics.com/single-cell-gene-expression/software/release-notes/build#header.

### CONTRIBUTIONS

For this study, KMB & QTE conceptualized the project. BFM, GBS, CLW, AVP, KM, BF, KH, VR, DP, TW, PP, KMT, & KMB developed methods for project analysis. JL (NIH) and KIK supported the recruitment of patients and collected data. QTE, ZC, DP, IS, MB, SMW, & KMB collected samples for analysis. QTE, BFM, GBS, CLW, AVP, BF, KH, VR, DP, KMT, JL (VCU) & KMB performed bioinformatic analysis. QTE, BFM, AH, & KMB performed additional data analysis. KMB & QTE wrote the original draft; QTE, BFM, CLW, AH, IS, KMT, JL (VCU), KIK, SAT, & KMB critically reviewed and edited the final manuscript.

## ACKNOWLEDGEMENTS

KMB firstly wants to acknowledge all the brilliant and generous teachers in the oral health research field he has had the privilege to learn from over the last 15 years. Our teams further want to acknowledge the tremendous efforts and support of the Human Cell Atlas, specifically Aviv Regev and Sarah A. Teichmann, for supporting the Oral & Craniofacial Bionetwork these past three years. Furthermore, we acknowledge that this article has become stronger and more comprehensive through early conversations with Steve Offenbacher, Scott Williams, Julie Marchesan, Karen Swanson, Adam Kurkiewicz, Vicky Morrison, Oliver Gibson, Justin Duplantis, and Peter Kharchenko. We further wanted to acknowledge the fantastic efforts of the NIH/NIDCR Dental Clinic, specifically Rachel Adam and Danielle Elangue, for their generous support to provide human tissues for testing and validation. This work was supported by generous start-up funds from the ADA Science & Research Institute (Volpe Research Scholar Award) and the Chan Zuckerberg Initiative/Foundation program Pediatric Networks for the Human Cell Atlas to KMB.

